# Methylomes reveal recent evolutionary changes in populations of two plant species

**DOI:** 10.1101/2024.09.30.615871

**Authors:** Kevin Korfmann, Andreas Zauchner, Bing Huo, Corinna Grünke, Yitong Wang, Aurélien Tellier, Ramesh Arunkumar

## Abstract

Plant DNA methylation changes occur hundreds to thousands of times faster than DNA mutations and can be transmitted transgenerationally, making them useful for studying population-scale patterns in clonal or selfing species. However, a state-of-the-art approach to use them for inferring population genetic processes and demographic histories is lacking. To address this, we compare evolutionary signatures extracted from CG methylomes and genomes in *Arabidopsis thaliana* and *Brachypodium distachyon*. While methylation variants (SMPs) are less effective than single nucleotide polymorphisms (SNPs) for identifying population differentiation in *A. thaliana*, they can classify phenotypically divergent *B. distachyon* subgroups that are otherwise genetically indistinguishable. The site frequency spectra generated using methylation sites from varied genomic locations and evolutionary conservation exhibit an excess of rare alleles. Nucleotide diversity estimates were three orders of magnitude higher for methylation variants than for SNPs in both species, driven by the higher epimutation rate. Correlations between SNPs and SMPs in nucleotide diversity and allele frequencies at gene exons are weak or absent in *A. thaliana*, possibly because the two sources of variation reflect evolutionary forces acting at different timescales. Linkage disequilibrium quickly decays within 100 bp for methylation variants in both plant species. Finally, we developed a deep learning-based demographic inference approach. We identified recent population expansions in *A. thaliana* and *B. distachyon* using methylation variants that were not identified when using SNPs. Our study demonstrates the unique evolutionary insights methylomes provide that SNPs alone cannot reveal.

## Background

Understanding how variation is partitioned within and between populations is a central goal in evolutionary biology. Increasing evidence suggests that epigenetic modifications can vary within and between populations (Richards, et al. 2017). Nevertheless, our knowledge of how epigenomes evolve through neutral and selective forces and, consequently, their capacity to reveal population demographic histories is limited. DNA cytosine methylation, a conserved epigenetic modification in eukaryotes (Feng, et al. 2010), occurs in plants within three sequence contexts: CG, CHG, and CHH (H represents A, T, or C) (Zhang, et al. 2018). DNA methylation is known to be involved in regulating gene expression, suppressing transposable element activity, and stabilizing the genome (Zhang, et al. 2018). Long-term changes in the methylome are mainly shaped by genomic alterations such as the proliferation of transposable elements and mutations affecting DNA methylation pathways, whereas heritable stochastic epimutations that occur independent of genomic background drive short-term changes (Vidalis, et al. 2016).

The unique characteristics of epimutations make them appealing markers for evolutionary investigations. In plants, CG methylation is maintained during cell division by METHYLTRANSFERASE 1 (MET1) (Zhang, et al. 2018). Spontaneous stochastic loss or gain of methylation at CG sites can be transmitted stably across multiple generations (Cortijo, et al. 2014). van der Graaf *et al*. have estimated the forward (gain of DNA methylation) and backward (loss of methylation) epimutation rates per CG site in the model plant *Arabidopsis thaliana* to be approximately 2.56×10⁻⁴ and 6.3×10⁻⁴, respectively, per generation per haploid methylome (van der Graaf, et al. 2015). These rates are several orders of magnitude higher than the mutation rate of 7×10⁻^9^ base substitutions per site per generation in the same population (Ossowski, et al. 2010). The increased rates of change at the epigenetic level may facilitate evolutionary inference over recent timescales, where the utility of slower-evolving genetic markers is limited (Yao, et al. 2021; Sellinger, et al. 2023).

As transiently induced epimutations can be mistaken for stably inherited markers, a recent study exposed genetically identical *A. thaliana* to various environmental conditions to identify genomic regions—termed clocklike regions—where CG divergence rates are independent of environmental stimuli (Yao, et al. 2023). While these regions are valuable for evolutionary studies, they are challenging to identify in most species. Therefore, the same study observed that gene body methylated regions (gbM) exhibit a pattern similar to that of CG divergence in clocklike regions (Yao, et al. 2023). gbM refers to a subclass of genes that display elevated CG methylation. They are mostly found in moderately and constitutively expressed housekeeping genes (Muyle, et al. 2022). As they can be identified based on the distribution of methylation in the gene bodies, their use may facilitate the investigation of evolutionary processes in nonmodel plant species.

Previous attempts to explore how methylation variants evolve have either neglected demographic (neutral) models, mixed environmentally sensitive and insensitive methylation sites, or did not adequately assess the interdependence of genetic and methylation variants. Charlesworth and Jain’s analytic framework for hypermutable sites (Charlesworth and Jain 2014) has been applied to create a methylation site frequency spectrum (mSFS) (Vidalis, et al. 2016). Previous studies generated mSFS for genic CG sites and the methylation states of gbM genes, finding either no selection or only weak selection acting on the methylome in *A. thaliana* (Vidalis, et al. 2016; Muyle, et al. 2021) and maize (Xu, et al. 2020). However, these studies do not incorporate an explicit demographic model while assessing selection. Another study found that positive selection acted on methylation variants linked to metabolite regulation in *A. thaliana*, even after accounting for demographic histories (Shirai, et al. 2021). Other studies in *Brachypodium distachyon* (Eichten, et al. 2020), *Fragaria vesca* (Le Vève, et al. 2024; Sammarco, et al. 2024), lettuce (Cao, et al. 2024), and grapevines (Rodriguez-Izquierdo, et al. 2024) have compared evolutionary inferences using genetic and methylation variants by analyzing all CG sites as the environmental sensitivity of individual sites is unknown in most plant species. Recently, Sellinger *et al*. utilized methylation as a secondary marker alongside SNPs to enhance demographic inference over short and long timescales (Sellinger, et al. 2023). The conceptual novelty was to consider that the underlying genealogy at a locus (*i.e.* the coalescent tree tracing back the ancestry of the sample back in time) is common for SNPs and SMPs (under a neutral model of evolution). This represents the first population genomics inference model combining DNA mutations, epimutations, and recombination along the genome, and the basis for developing a Sequential Markovian Coalescent method (SMCm). However, that study exhibits a fair amount of noise in the inference of past evolutionary history when it was applied to methylome data from *A. thaliana* [8]. These inaccuracies are due to a lack of knowledge about the optimal SMP markers to use, the relevant genomic context, the relationship between SNPs and SMPs, and the SMCm method being very sensitive and not accounting well for methylated region effects (so-called differentially methylated regions or DMRs). A related question is whether DMRs can be used for evolutionary inferences as well.

Given these limitations, our goal was to assess how well methylation variants reveal known evolutionary patterns inferred using SNPs. We chose *A. thaliana* and *B. distachyon* for our analyses because their evolutionary histories are well-described (Kawakatsu, et al. 2016; Eichten, et al. 2020). We found that the capacity of methylation variants to differentiate populations is contingent on when populations diverge. We also show that CG sites across the methylome capture within- and between-population patterns comparable to environmentally insensitive sites (clocklike regions and gbM) (Yao, et al. 2023) in samples grown in a common garden. We found weak or no local correlations between genetic and methylation variants, indicating they reflect different aspects of population evolutionary histories. Additionally, the linkage between neighboring methylation variants was minimal. Conventional genome-based demographic inference methods, such as the Sequential Markovian Coalescent (Strütt, et al. 2023), show limitations in using methylation variants due to high sensitivity to violations of the Markovian properties of the underlying coalescent model along the genome [8]. Moreover, since such methods require knowing the variant accumulation rates and distributions—often unknown for methylation in many species—we developed a deep learning method that uses methylation as a neutral marker to estimate past demography. Unlike conventional convolutional networks, we use (to our knowledge) the first Encoder-Only Transformer architecture (Vaswani, et al. 2017; Devlin, et al. 2018; Korfmann, et al. 2023) method, focusing on the placement of mutations on a hidden ancestral recombination graph. We track estimates of coalescent events in a time-discretized transition matrix by using heterogeneous sites per genomic window as input, thereby applying a coarse-grained approach, similar to tsABC (Strütt, et al. 2023). By aligning training and real data distributions, we show that rapidly changing methylation variants can reveal recent demographic shifts in selfing and clonal species.

## Results

### CG methylation variants identify population structure in recently diverged groups

We analyzed genetic and methylation diversity using 63 samples each from the Central European (CEU) and the Iberian non-relicts (IBnr) *A. thaliana* populations (Table S1). These populations were used by (Strütt, et al. 2023). Across the two populations, 31-33% of CG methylation sites were polymorphic (Figure S1A), which is similar to a previous estimate for CG sites in *A. thaliana* mutation accumulation lines (Becker, et al. 2011). In contrast, 1.6-1.7% of coding sites and 2.7-2.8% of non-coding sites were polymorphic (Figure S1A). Interestingly, 80–81% of sites in clocklike regions and 60% of sites in gbM regions were polymorphic, which is significantly higher compared to CG sites found elsewhere in the genome (Figure S1A).

We compared our results from *A. thaliana* with those from 18 samples from the clonally propagating grass *B. distachyon*. Wilson *et al*. used SNPs to identify clonal groups among Turkish *B. distachyon*, with samples within each clonal group showing high levels of genetic similarity (Wilson, et al. 2019). Eichten *et al*. identified high levels of consistency between the known genetic relationships for the clonal groups and the extent of CG divergence (Eichten, et al. 2020). Through the analysis of a larger number of Turkish accessions, Skalska *et al*. identified two distinct genetic clusters: one in coastal regions and the other in central regions, the latter roughly corresponding to the high-elevation Anatolian plateau (Skalska, et al. 2020). In this study, we focus exclusively on a single clonal group, as defined by Eichten *et al*. (clonal group 1), and analyze only the 18 samples from the central Turkish region that have average bisulfite sequencing depths greater than 10 (Eichten, et al. 2020) (Table S1). Between 0.8-2.8% of CG methylation sites were polymorphic in *B. distachyon* (Figure S1B). Similar to *A. thaliana*, the frequency of segregating CG methylation sites was higher within gbM regions, with 3–4% of sites segregating (Figure S1B). The median length of DMRs in *B. distachyon* was 107 bp (1^st^ percentile: 44 bp, 99% percentile: 717 bp) which is smaller than the median length in *A. thaliana* DMRs, which was 181 bp (1^st^ percentile: 78 bp, 99% percentile: 1125 bp) (Figure S2).

We estimated the proportion of sites at which two samples were different (p-distance) and clustering to uncover how the net total of genetic and methylation variation differed between samples from the CEU and IBnr populations. In this approach, a clear separation between the CEU and IBnr populations was apparent with both coding and non-coding single nucleotide polymorphisms (SNP) (Figure 1A). Although single methylation polymorphisms (SMP) and differentially methylated regions (DMR) largely separated the two populations, some samples were separated by only minimal distances (Figure 1A). Clustering indicated nine or more groups within the SNP data (Figure S3A-B), whereas only two groups were apparent in the SMP and DMR datasets (Figure S3C-D).

**Figure 1.**
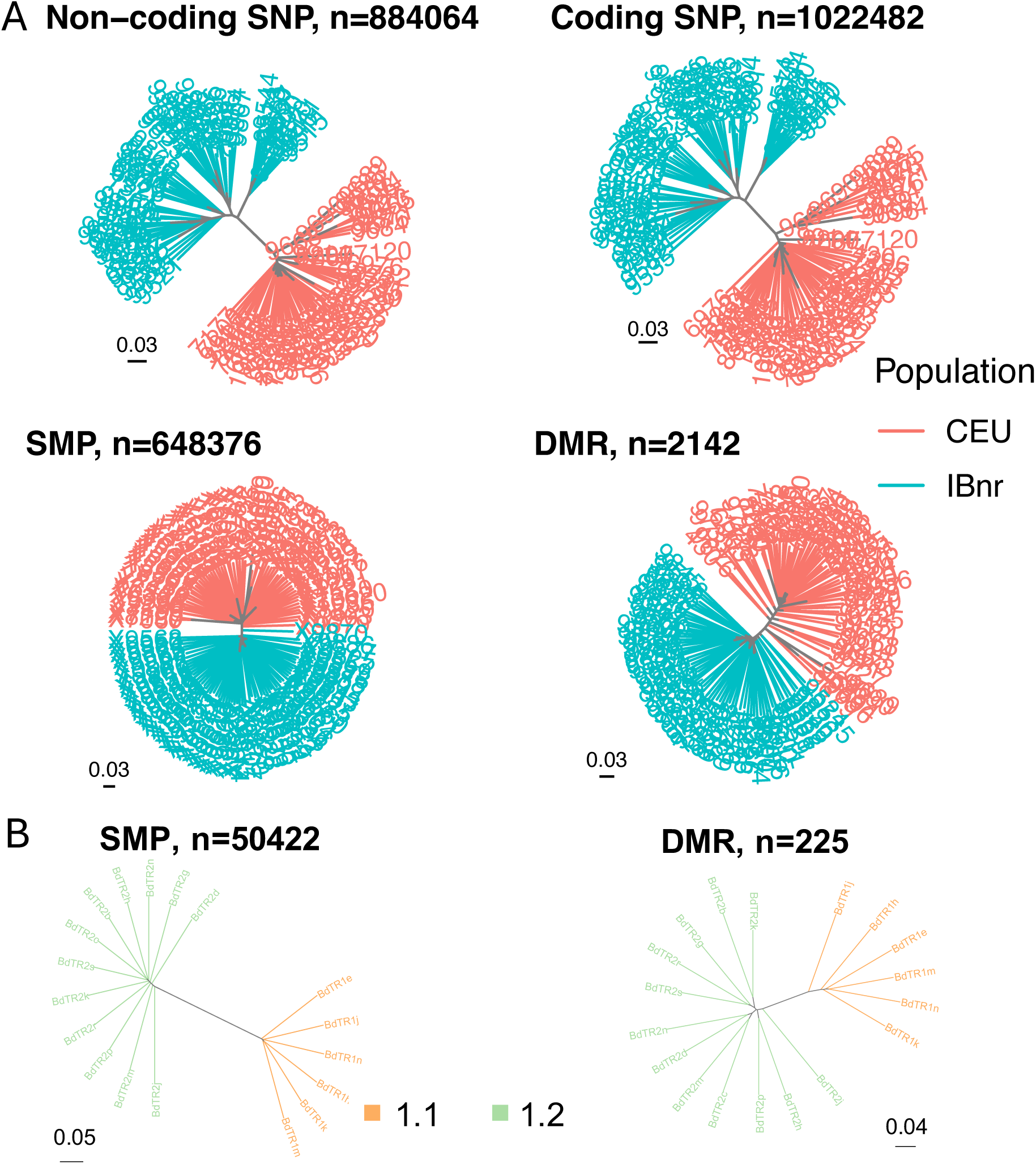
Methylomes outperform single nucleotide polymorphisms (SNP) in detecting population subdivision in *Brachypodium distachyon* but not *Arabidopsis thaliana*. (**A**) Neighbor-joining trees were generated using p-distance matrices for non-coding SNPs, coding SNPs, clocklike region single methylation polymorphisms (SMP), and clocklike region differentially methylated regions (DMR) for Central European (CEU) and the Iberian non-relicts (IBnr) *A. thaliana*. (**B**) Neighbor-joining trees generated using p-distance matrices estimated for SMP and DMR in gbM regions of a Turkish *Brachypodium distachyon* clonal group.

Furthermore, we performed discriminant analysis of principal components (DAPC) analyses to identify how correlated changes in subsets of genetic and methylation variants can be used to discriminate the two populations. DAPC indicated greater discriminatory power with coding and non-coding SNPs compared to SMPs and DMRs (Figure S3E). Although p-distance matrix-based comparisons could have been influenced by the greater number of SNPs (Figure 1A), marker numbers were comparable for the DAPC, and yet SMPs and DMRs showed poor discriminatory capability (Figure S3E). SMPs and DMRs that occur within gbM genes have comparable performance to those occurring within clocklike regions in discriminating the two populations (Figure S4). High methylation rates in *A. thaliana* may have caused multiple changes at the same site (so-called homoplasy accounted for as a finite site model (Charlesworth and Jain 2014; Wang and Fan 2014; Vidalis, et al. 2016)), resulting in the underestimation of the true levels of divergence. Indeed, the divergence between IBnr and CEU populations is estimated to have occurred after the last glacial maximum ca. 10,000-20,000 years ago (Hewitt 1999; Sharbel, et al. 2000; The1001GenomesConsortium 2016).

We then performed a similar analysis using the *B. distachyon* accessions. Unlike the *A. thaliana* populations, there were just 63 segregating SNP sites within the *B. distachyon* clonal group 1, suggesting a very recent origin for this genetically homogeneous clone. We expected that methylomes would be more useful in revealing population structure during the very early stages of clonal expansion and divergence. Even though both samples were genetically identical (Wilson, et al. 2019) and originated from a similar elevational gradient in the central Turkish region, SMPs and DMRs revealed two distinct subgroups (Figures 1B, S5). Both our results using gbM regions and the results from Eichten *et al*. that use the entire collection of CG sites (Eichten, et al. 2020) reveal a difference in methylation profiles between the two subgroups of Group 1. The SMP and DMR based trees were longer compared to those in *A. thaliana* (Figure S4A) indicating greater methylome differentiation in *B. distachyon*. We obtained phenotype data for the two subgroups (Filiz, et al. 2009) and found significant differences in the longitude, seed lengths, seed yield, seed production, and germination percentages (Figure S6-S7). As expected, we did not find any significant difference in elevation between the subgroups from the Central Turkish region (Figure S7). Our findings suggest that methylation variants can reveal subpopulation structure in recent timescales, where there has been little time for genetic differences and multiple methylation changes to accumulate. However, they become saturated with multiple changes occurring at the same site as time passes obscuring population differentiation.

We sought to understand how genomic contexts influenced the methylation site frequency spectra (mSFS) in *A. thaliana*. The mSFS reveals how methylation rates, demographic changes, and natural selection influence changes in the methylome over time (Charlesworth and Jain 2014; Vidalis, et al. 2016). When considering all CG SMP or DMRs jointly, we found a greater frequency of unmethylated alleles for the CEU and IBnr populations, but IBnr consistently had higher frequencies of unmethylated alleles for SMP sites at all genomic regions (Figure 2A). The overall shapes of the mSFS for SMPs are consistent with previous findings that the backward methylation rate promoting the change from methylated to unmethylated is higher than the forward rate (van der Graaf, et al. 2015) and are consistent with previous mSFS generated by combining *A. thaliana* samples from extended geographic regions (Vidalis, et al. 2016; Muyle, et al. 2021).

**Figure 2.**
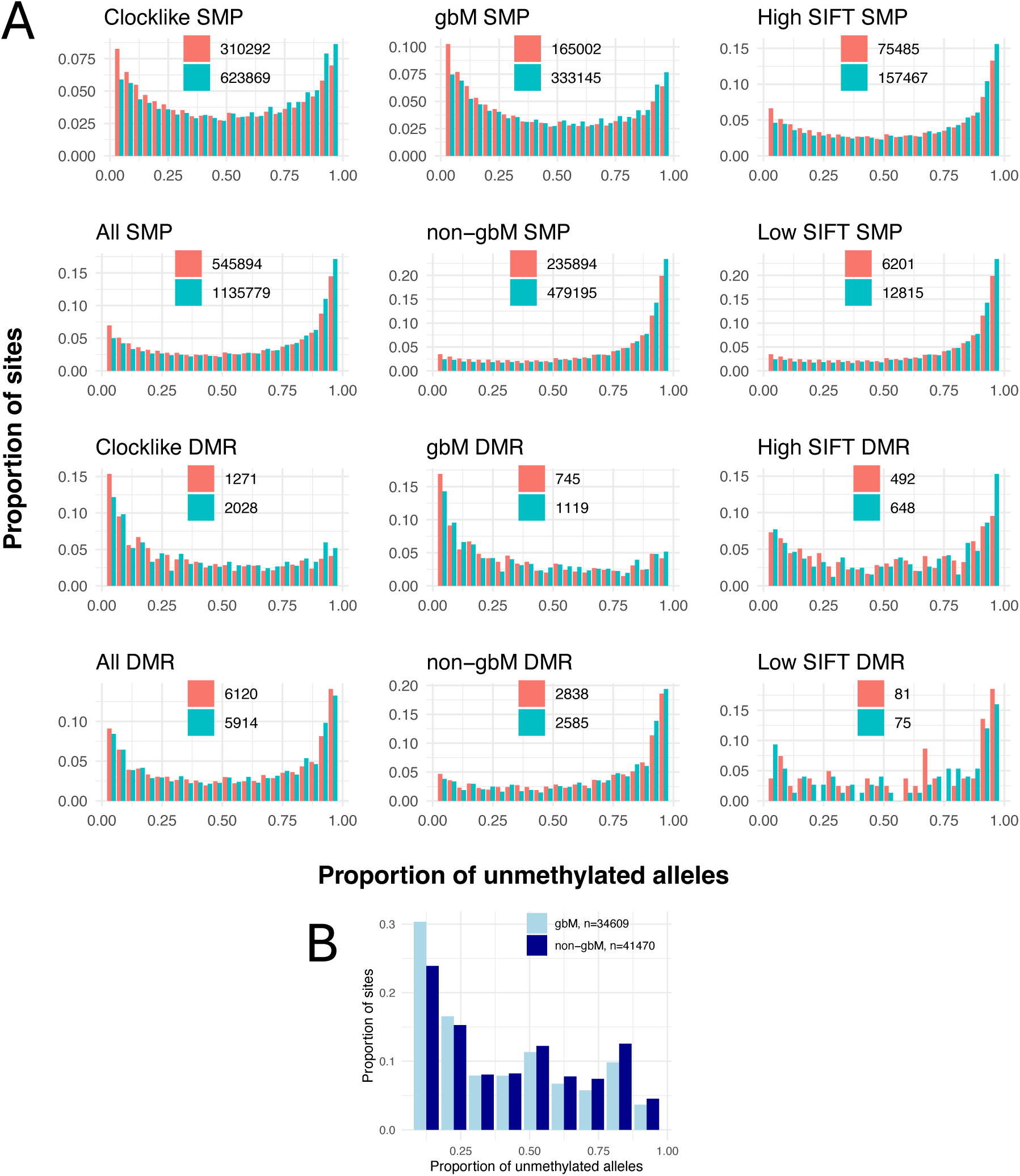
Methylation site frequency spectra generated using single methylation polymorphisms were similar regardless of genomic contexts. Methylation site frequency spectra (mSFS) generated using differentially methylated regions (DMR) exhibited greater variability. (**A**) mSFS for the Central European (CEU, Salmon) and the Iberian non-relicts (IBnr, Cyan) *Arabidopsis thaliana* populations using CG sites that occur in clocklike, gene body methylated (gbM), non-gbM, and low and high Sorting Intolerant from Tolerant (SIFT) score regions. Only sites with minor allele frequencies (MAF) >0.02 were used to generate the mSFS. (**B**) mSFS for *Brachypodium distachyon* clonal group 1.2 for single methylation polymorphisms (SMP) and DMRs occurring within gbM and non-gbM regions. Only polymorphic sites were used to generate the mSFS. For (A) and (B), the numbers in each panel indicate the number of sites used for the analyses.

The relative frequencies of methylated and unmethylated alleles varied across individual genomic regions. SMPs in gbM regions for CEU and DMRs in gbM regions in both populations showed an excess of methylated alleles (Figure 2A). gbM are defined based on an excess of CG methylation and this is reflected in the mSFS. The number of methylated and unmethylated alleles was similar for SMP sites in clocklike regions, which could partly result from their comparable forward and backward methylation rates (Yao, et al. 2023). However, the differing direction of the skew in CEU and IBnr indicates some differences in the methylation rates for the populations for these sites. We also studied the mSFS for intervals containing genes with low and high Sorting Intolerant From Tolerant (SIFT) scores, which are positively correlated with the level of evolutionary constraint that genes are under (Ng and Henikoff 2003). Our results indicate that the level of gene constraint has little impact on the mSFS shape (Figure 2A). While individual genomic regions differ in the proportion of methylated sites they contain, it is evident that all cases exhibit an excess of rare variants, suggesting a shared influence of demographic changes on all classes of variants.

In contrast to the mSFS for SMPs in *A. thaliana* (Figure 2A), the mSFS of *B. distachyon* group 1.2 indicated a slight excess of methylated alleles within gbM and non-gbM SMPs, although it was less pronounced for the latter (Figure 2B). Similar to *A. thaliana*, an excess of rare alleles was observed, suggesting a common demographic influence.

We next sought to understand how SNPs and methylation polymorphisms are distributed within *A. thaliana* populations. We first calculated the pairwise nucleotide diversity (π) for SMPs (π*_SMP_*) and SNPs (π*_SNP_*) from noncoding, clocklike, gbM, non-gbM, and high and low SIFT genic regions. We scaled estimates by the total number of sites (polymorphic and monomorphic) to ensure that the results were comparable (that is per-site estimates). π*_SMP_* is three orders of magnitude higher than π*_SNP_* (Figure 3A), likely driven by the high rates of change in methylomes (van der Graaf, et al. 2015). π*_SNP_* in the CEU population and π*_SMP_* in CEU and IBnr populations for clocklike regions were significantly higher than in other regions (Figure 3A). This may indicate improved detection of environmental insensitivity—the criterion for defining clocklike regions—when more polymorphic sites are present. π*_SMP_* was similar for gbM and non-gbM regions indicating that the levels of polymorphisms are not affected by gbM (Figure 3A). π*_SMP_* was also similar for regions containing genes with high-SIFT scores or low-SIFT scores (Figure 3A) indicating that constraint at the genic level has negligible effects on the levels of local methylation diversity.

**Figure 3.**
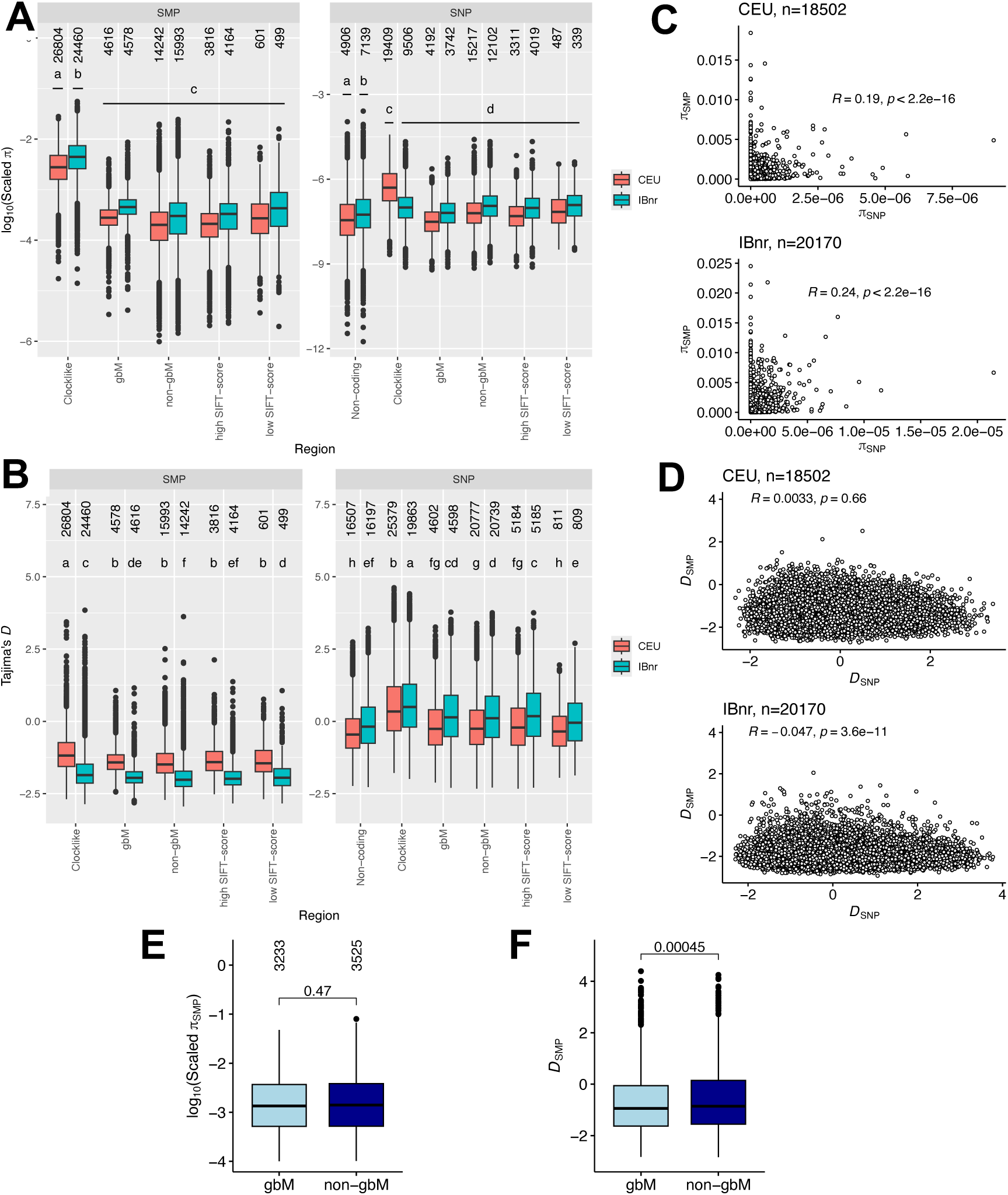
Unique evolutionary signatures in genomes and methylomes. (**A**) Estimates of pairwise nucleotide diversity (π) scaled by the number of cytosines per region for single methylation polymorphisms (π*_SMP_*) and the number of bases for single nucleotide polymorphisms (π*_SNP_*) for the Central European (CEU) and the Iberian non-relicts (IBnr) *Arabidopsis thaliana* populations. (**B**) Estimates of Tajima’s *D* per region for CEU and IBnr. Comparisons of (**C**) π*_SNP_* versus π*_SMP_* and (**D**) Tajima’s *D* for SNPs (*D_SNP_*) and SMPs (*D_SMP_*) per gene for CEU and IBnr. The numbers in the titles of each panel indicate the number of sites used for the analyses. Correlation tests were conducted to assess the associations between π and Tajima’s *D*. (**E**) π*_SMP_* scaled by the number of cytosines per region for *Brachypodium distachyon* clonal group 1.2. (**F**) *D_SMP_* per region for *B. distachyon* group 1.2. For (A), (B), (E), and (F), the numbers above the boxplots indicate the number of intervals. For (A) and (B), letters indicate group-wise differences in means following Tukey’s honest significant difference tests. For (E) and (F), *P*-values from Wilcoxon tests are shown. For (F), the color scheme and number of sites are the same as in (E).

We then compared how selective and demographic processes influence genomes and methylomes by estimating Tajima’s *D* for CEU and IBnr. For methylomes, we used a modified Tajima’s *D* using the *D^m^* (Wang and Fan 2014) that accounts for the hypermutability of the methylation sites. *D_SNP_* was slightly negative for noncoding sites in the CEU and IBnr populations (Figure 3B), reflecting the known history of population expansions (Strütt, et al. 2023). The higher *D_SNP_* for IBnr (Figure 3B) is consistent with larger effective population sizes over recent timescales (Strütt, et al. 2023). In contrast, IBnr had significantly lower *D_SMP_* compared to CEU (Figure 3B). This could be due to variation in methylation rates, purifying selection, or demographic and selective (purifying selection) histories. Therefore, we used SMCm (Sellinger, et al. 2023), which employs fixed mutation rates and compares the distributions of mutation and methylation markers, to estimate the forward and backward methylation rates for the IBnr and CEU populations. Methylation rates were only slightly higher in CEU but the ratio of backward to forward rates was similar in both populations (Figure S8). Hypermutability should result in greater frequencies of intermediate variants (Charlesworth and Jain 2014), but we did not observe such a pattern (Figure 2A). Tajima’s *D* is based on a comparison of the number of segregating sites versus π. The disparity in *D_SMP_* appears to be driven primarily by the variation in the number of segregating sites per region (Figure S9). This difference suggests that the demographic histories of the two populations may be slightly different.

We then estimated Tajima’s *D* in different genomic contexts for CEU and IBnr. Clocklike regions had significantly higher *D_SNP_* and *D_SMP_* compared to other regions (Figure 3B). The *D^m^* method assumes that forward and backward methylation rates are equivalent (Wang and Fan 2014). Although the ratio of the backward to forward methylation rates in clocklike regions is 1.36, the ratios are 2.11 and 2.84 for gbM and non-clocklike regions, respectively (Yao, et al. 2023), indicating the assumption of *D^m^* is less strongly violated for clocklike regions. The differences in *D_SMP_* between high-SIFT vs low-SIFT or between gbM and non-gbM were not consistent in both populations (Figure 3B). This contrasts with consistently lower *D_SNP_* for regions containing genes with low-SIFT score that are under lower selective constraints compared to regions with genes with higher-SIFT regions (Figure 3B). While ignoring the differences between regions can introduce some noise when inferring evolutionary signatures from methylomes, our results suggest any such effect is minimal. There was only a weak positive correlation for π (Figure 3C) generated for SNPs vs SMPs that occur in coding regions and negligible correlation for Tajima’s *D* (Figure 3D).

We compared π and Tajima’s *D* for *B. distachyon* SMPs generated using SNPs for the Anatolian and Turkish lineage (B_East) (Stritt, et al. 2022; Minadakis, et al. 2023), to which clonal group 1.2 belongs. Scaled π for SMPs was larger for the clonal group 1.2 compared to the *A. thaliana* populations, but there were no consistent differences between gbM and non-gbM regions (Figure 3A, 3E). Similar to *A. thaliana* (Figure 3A), scaled π for SMPs for clonal group 1.2 was three orders of magnitude larger than the median SNP-based scaled π (∼1.8×10⁻⁶) for the B_East lineage (Stritt, et al. 2022; Minadakis, et al. 2023) (Figure 3E). Similar to *A. thaliana* (Figure 3B), *D_SMP_* for clonal group 1.2 was negative. *D_SMP_* was slightly but significantly larger in non-gbM regions compared to gbM regions (Figure 3F). These negative values contrast with the positive *D_SNP_* for SNPs in the B_East lineage (Minadakis, et al. 2023), further suggesting that methylomes exhibit unique signatures.

An important consideration for population genetic analysis is the presence of sites with independent evolutionary histories (Sellinger, et al. 2023). Therefore, we assessed the levels of pairwise linkage disequilibrium (LD) to identify the number of independent polymorphic SNPs and SMPs. Pairwise LD decay was quicker for SMPs compared to SNPs in *A. thaliana* (Figure 4A). LD decay was much stronger for SMPs occurring within clocklike or gbM regions than for coding and noncoding SNPs with r^2^ dropping below 0.1 within the first 100 bp (Figure 4A). LD decays to 0.1 for SMPs sites within gbM regions for *B. distachyon* within the first 100 bp (Figure 4B).

**Figure 4.**
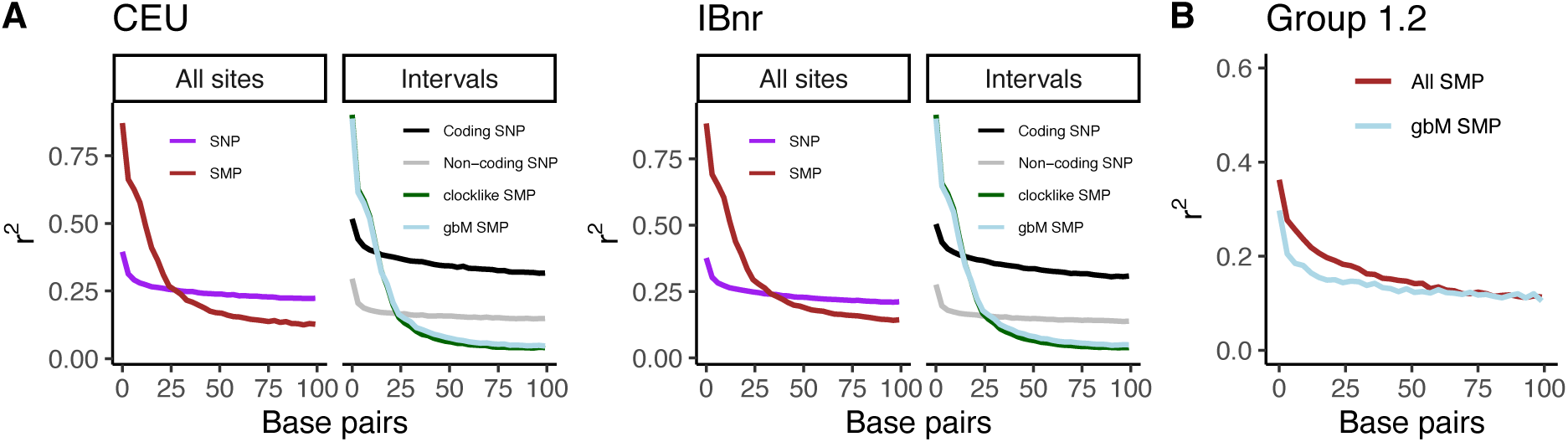
Weak linkage disequilibrium (LD) between neighboring single methylation polymorphisms (SMP). (**A**) Pairwise r^2^ estimated using all single nucleotide polymorphisms (SNP), coding SNPs, non-coding SNPs, all SMPs, and SMPs occurring within clocklike regions, SMPs occurring within gene body methylated regions (gbM) for Central European (CEU) and Iberian non-relicts (IBnr) *Arabidopsis thaliana* populations. (**B**) Pairwise r^2^ was estimated using SMPs for *Brachypodium distachyon* clonal group 1.2.

### Demographic inference using genetic and methylation variants

We developed a novel inference method that focuses on large genome blocks (20 Kbp windows) likely spanning multiple coalescent genealogies (Figure 5A), to estimate the times to the most recent common ancestor (TMRCA). Our method assumes that the number of heterozygous markers per window is correlated with the average TMRCA. Essentially, longer coalescent times allow more mutations and epimutations to accumulate. To mitigate the possibility that older genealogies (long TMRCA) can be unexpectedly short due to immediate recombination, our approach uses 20 Kbp windows to average out genealogies. We apply a scaling to the number of markers to estimate the TMRCA (Figure S10). Although the chosen scaling factor will not perfectly offset all variable demographic samples—given that we trained using a fixed recombination rate and a fixed window size—offsets are minimal and addressed during training. An encoder-only transformer neural network processes this data, learns the scaling factor, and predicts population sizes over time. This is done by summarizing the TMRCA of two consecutive windows in the empirical transition matrix. The differences between each pair of samples are used to infer their TMRCA. The TMRCA values for all pairs across multiple intervals (windows) in the genome are then represented as a transition matrix. This method combines raw genotype data restructuring with deep learning to infer demographic history.

**Figure 5.**
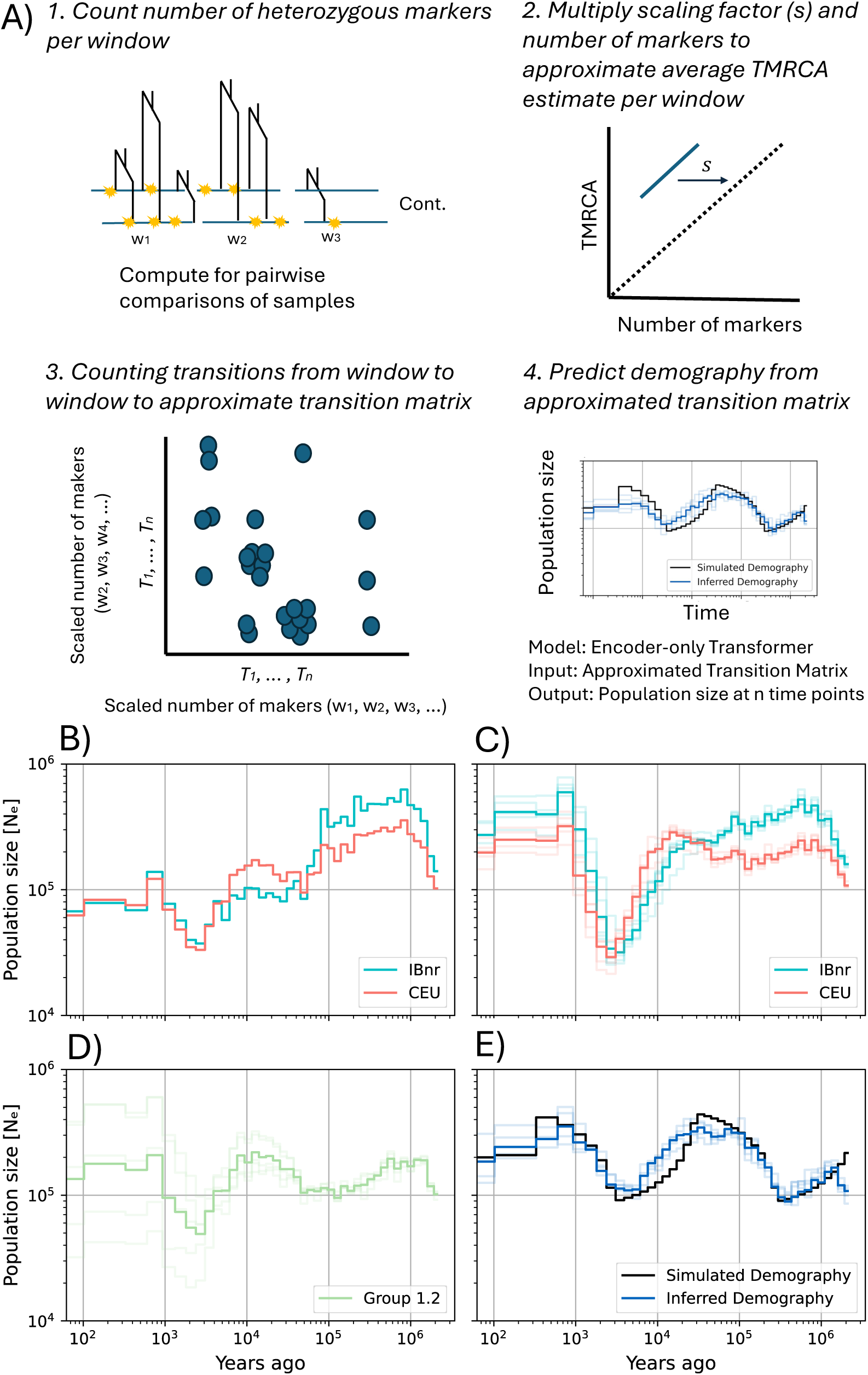
Demographic inference using mutation or methylation markers. (**A**) Schematic of the approximated transition matrix inference framework. The genome is first divided into 20 Kbp bins, and heterozygous markers are counted for all pairwise comparisons in a sample of 10 individuals. These counts are then scaled by an empirically chosen factor to ensure sufficient values in the approximated transition matrix by aligning the number of markers per window to the TMRCA in that window. The framework constructs this matrix by concatenating all pairwise comparisons into a long sequence and applying a sliding window of size 2 and step size 1. Each pair of consecutive values in this window represents coordinates in the transition matrix, which is incremented at those positions. This process results in an unnormalized transition matrix that captures the genetic relationships between samples. Finally, an encoder-only transformer neural network takes this noisy matrix as input, implicitly learning the fixed scaling factor, and predicts population sizes at various time points. Inference using over 42 time steps using (**B**) single nucleotide polymorphisms (SNP) and (**C**) single methylation polymorphisms (SMP) for the central European (CEU) and the Iberian non-relicts (IBnr) *Arabidopsis thaliana* populations. (**D**) Demographic inference using SMP for *Brachypodium distachyon* groups 1.2. SMP inference was done using sequences from all five chromosomes (light lines, average in bold), while SNP inference is based on chromosome one. (**E**) Inference of simulated sawtooth demography using five replicates, 4Mbp segments, and 10 samples.

For deep learning inference, the distributions of heterozygous markers in training and real data must overlap to ensure in-distribution inference. For the training data, we collapsed each training transition matrix into a single scalar by summing the flattened matrix and plotted the distribution of unnormalized transition matrices (Figure S11A). To ensure the real data *from A. thaliana* and *B. distachyon* matched the training data, we analyzed 20 randomly sampled genomic regions. For the SNP data, we selected start positions uniformly between 0-28 Mbp, whereas for the methylation data, the start positions were chosen uniformly between 0-0.5 Mbp. For both data types, we determined each region’s length by sampling from a uniform distribution between 1-3 Mbp. Summing the resulting transition matrices for the IBnr, CEU, and Group 1.2 regions (Figure S11B-D) produced values within the training data range (Figure S11A). Despite the differential properties of SMPs and SNPs, the distribution of heterozygous markers per 20 Kbp window was similar across populations (Figure S11B-C), supporting the use of a common training dataset for deep learning.

We applied our coarse−grained inference approach to *A. thaliana* and *B. distachyon*. We generated approximated transition matrices from 10 samples and 4 Mbp sequence lengths. The 45 pairwise comparisons totaling 180 Mbp with 20 Kbp windows yielded 9,000 pairs of TMRCA values per matrix. We detected a population bottleneck ∼100,000 years ago (100 kya) in CEU and IBnr using SNPs (Figure 5B), which was consistent with the findings of a previous study (Strütt, et al. 2023). This signal was not clearly detected with SMPs (Figure 5C), indicating their limited use for ancient events. Both SNPs and SMPs indicated another bottleneck between 1-10 kya (Figure 5B-C), but SMPs also revealed a subsequent population expansion (Figure 5C) undetected by SNPs (Figure 5B). The steeper expansion of IBnr compared to CEU (*A. thaliana*) in the last 10,000 years likely led to the observed excess of rare methylation alleles (Figures 2A, 3B). Using SMPs, we recovered two bottlenecks (∼100 kya and between 1-10 kya) in *B. distachyon* (Figure 5D).

We validated our demographic inference method with two approaches. First, we simulated the sawtooth demography observed in both species, confirming that our deep learning method accurately detected multiple bottleneck and expansion events (Figure 5E). Second, we simulated methylation markers with high rates of change (10⁻⁴, 10⁻⁵, or 10⁻⁶ per site per generation) and backward rates 2.5 times larger than forward rates, based on the estimates provided in (van der Graaf, et al. 2015), to test whether deviations from the infinite sites model affect inference. We found that our method accurately recovered demographic histories beyond 0.6 kya and before 5, 22, and 442 kya for methylation rates of 10⁻⁴, 10⁻⁵, and 10⁻⁶ per site per generation, respectively (Figure S12). Conservatively, this suggests that inferences 20 kya and beyond are unlikely to be accurate for CEU and IBnr due to their estimated rates of methylation change being around 10⁻⁵ per site per generation (Figure S8). Note that the demographic estimates with SNPs are likely accurate for older events (>20kya).

Finally, we obtained short tandem repeat (STR) calls from (Reinar, et al. 2021) to test whether this independent, fast-mutating marker also provides evidence for population expansions in the *A. thaliana* populations. STRs occurring within noncoding regions were obtained from 36 and 54 samples from the CEU and IBnr, respectively, that overlapped with our methylation dataset. We used the Bottleneck software (Cornuet and Luikart 1996) to test for a heterozygosity deficit, which can serve as evidence of a population expansion (Luikart and Cornuet 1998; Girod, et al. 2011). Since we were working with highly inbred effectively haploid samples, we combined the calls from two randomly chosen pairs of samples to create diploid data. We consistently detected a significant heterozygosity deficit using the sign test, the standardized difference test, and the Wilcoxon signed-rank test in CEU and IBnr (Table S2). These results indicate that we are confidently estimating recent population expansion using SMPs (Figure 5C), highlighting the potential of methylation markers to provide insights beyond those offered by SNPs alone.

## Discussion

We observed contrasting patterns in the capacity of methylation variants to distinguish population subdivisions in *A. thaliana* and *B. distachyon*. In *A. thaliana*, the infinite sites model is likely violated for methylation, resulting in the erosion of methylome differentiation likely due to the older divergence between populations. In contrast, genetically identical *B. distachyon* samples exhibited distinct methylation profiles, most likely driven by their very recent divergence. While individual genomic locations varied in whether they had more methylated or unmethylated alleles—likely driven by the underlying molecular architecture—CG methylation sites and DMRs consistently showed an excess of rare alleles in *A. thaliana* and *B. distachyon*, indicating a common demographic influence. We found weak negative correlations between π and no local correlations for Tajima’s *D* between SNPs and methylation variants in *A. thaliana*, underscoring their distinct molecular and evolutionary properties, similar to the conclusions reached by (Schmitz, et al. 2013). Linkage quickly decays between neighboring SMPs, which was also seen in wild *A. thaliana* sampled across the Northern Hemisphere (Schmitz, et al. 2013). SMPs and SNPs recover recent and older demographic changes, respectively, providing a comprehensive reconstruction of population evolutionary histories.

The differing ability to detect population subdivisions in *A. thaliana* and *B. distachyon* using methylation variants could lead to differences in divergence times, methylation rates, or local adaptation. As the post-glacial re-colonization of the Iberian Peninsula and the spread northward started ∼16 kya (Hewitt 1999), multiple changes at the same methylation site (homoplasy) during this period may have caused an underestimation of the true levels of divergence in *A. thaliana*. The lack of congruence between SMP- and SNP-based comparisons in A. thaliana also suggests that there has been limited post-expansion gene flow, as otherwise, the two profiles would be more similar. The greater capacity to detect group subdivision is *B. distachyon* could reflect a recent separation by longitude contributing to methylome divergence. We indeed rule out the role of adaptation such as found in *F. vesca* (Le Vève, et al. 2024), because we generated inferences using gbM regions that are highly conserved in angiosperms (Muyle, et al. 2022). The observed divergence could be due to low rates of methylation change, but we could not estimate methylation rates for *B. distachyon* as our rate estimate relies on comparing the distributions of SNP and methylation variants. The most parsimonious explanation is that the two subgroups diverged recently. As the B_East lineage split from the western Mediterranean lineage 23 kya (Stritt, et al. 2022; Minadakis, et al. 2023), the split between the closely related subgroups is likely to have been far more recent, ensuring that the infinite sites model was not violated for methylation variants. Our findings underscore the necessity for methods distinguishing neutral demographic changes from selective pressures on the methylome, elucidating why SMPs behave differently across species.

The differences between the shapes of the mSFS for the different genomic contexts indicate that potential for differences in their molecular basis. gbM results from the combined activity of MET1, which is enhanced by nearby existing methylation, as well as the histone variant H2A.Z and the DNA demethylase ROS1 (Briffa, et al. 2023). The molecular basis underlying the clocklike, non-gbM regions, as well as regions with high and low SIFT scores, remains unknown. Interestingly, the shapes of the mSFS from SMPs in clocklike regions vary between IBnr and CEU, and both also differ from the mSFS of DMRs in the same regionsThis pattern, combined with their much higher rates of polymorphism compared to the remaining genomic regions, suggests that these intervals likely have a unique molecular basis that warrants further research. As clocklike regions accumulate methylation differences in a linear fashion over time, they can be useful for studying neutral signatures (Yao, et al. 2023). While gbM sites may not be entirely neutral—partly because they are known to affect gene expression (Muyle, et al. 2022)—the similarity of their mSFS to those of clocklike regions, along with the fact that ∼60% of clocklike regions occur within gbM regions (Yao, et al. 2023), suggests shared molecular features and that gbM sites may serve as a suitable substitute when clocklike regions have not been defined.

Our results support the use of individual CG sites from all genomic contexts for population-genetic analyses. Our between-population comparisons using CG sites in gbM regions successfully identified the known population structure within *B. distachyon*, similar to a previous study using genic CG sites (Eichten, et al. 2020). Sites from various genomic contexts showed variation in the prevalence of methylated versus unmethylated alleles. However, a consistent feature was the excess of rare alleles across these sites. Given the rapid changes in markers such as methylation, determining the ancestral state is challenging. The skew toward rare alleles in the mSFS suggests that demographic parameters exert a consistent impact, even when the molecular mechanisms controlling different genomic regions vary. Moreover, the *A. thaliana* (Kawakatsu, et al. 2016) and *B. distachyon* (Eichten, et al. 2020) methylomes, generated under similar environmental conditions, ensure that sites affected solely by transient environmental effects would not be recognized as SMPs. Another study using all CG sites from lettuce methylomes, cultivated under uniform environmental conditions, also demonstrated the capacity to distinguish multiple species (Cao, et al. 2024). DMRs may be preferable when low sequencing depth limits SMP identification (Taudt, et al. 2018), but their unclear evolutionary basis and susceptibility to multiple forces (Sellinger, et al. 2023) make them unreliable when used on their own.

The weak local correlations between genetic and methylation diversity and the association between gene constraint and methylation diversity in *A. thaliana* can partly be attributed to methylation affecting the mutability of neighboring sites (Mugal and Ellegren 2011; Xia, et al. 2012; Kusmartsev, et al. 2020) and transposable element polymorphisms influencing methylation variation (Stuart, et al. 2016; Shen, et al. 2018). However, only about 12% of methylation quantitative trait loci (QTLs) in *A. thaliana* are within 10Kbp of the epiallelic target (Schmitz, et al. 2013). In maize, approximately 50% of methylation QTLs occur in *cis* (Eichten, et al. 2013), indicating a potentially stronger trend in this species. The higher rate of methylation changes compared to mutation accumulation (Ossowski, et al. 2010; van der Graaf, et al. 2015) may also explain why methylation differences persist despite genetic homogeneity. Our simulations also show that coalescent histories of markers with vastly different rates of change diverge, which can explain the general lack of correlation between π and Tajima’s *D* for local genetic and methylation diversity. Our results are similar to those of previous studies in lettuce (Cao, et al. 2024) and cotton (Zhao, et al. 2024), which also found higher levels of polymorphism in the methylome compared to the genome. The lack of correlation or similarity between the Tajima’s *D* estimates from the genome and methylome in *A. thaliana* and *B. distachyon* supports the notion that the SNPs and SMPs likely reflect different parts of the common underlying coalescent genealogy at a locus: SNPs reflect the older part of the tree, whereas SMPs would reflect the topology and branch lengths in the more recent past. It is then not surprising that for recent positive selection events (selective sweeps), the signatures are similar for both types of markers, and SMPs may even be more accurate in detecting them (Shirai, et al. 2021). Our study mostly focuses on local correlations in diversity, rather than genome-wide correlations, where life history traits and genetic differentiation levels also play a significant role (Lele, et al. 2018; Medrano, et al. 2020; Wang, et al. 2020; Avramidou, et al. 2021; Fargeot, et al. 2021; Avramidou, et al. 2023; Langford, et al. 2024).

Our study demonstrates that methylation variants can independently detect and allow inference of historical demographic changes. This represents a significant improvement over a previous approach (Sellinger, et al. 2023), where methylation was used only as a secondary marker to supplement SNP-based inferences. Furthermore, our improved deep learning method does not require precise methylation rate parameters and is less sensitive to the presence of invariant methylation sites (compared to the SMCm in (Sellinger, et al. 2023)). Instead, it trains on diverse scenarios to match observed variant distributions. Our simulations confirmed the utility of rapidly mutating markers for inferring demographic changes. Our STR analysis confirm that there have been recent population expansions in the two *A. thaliana* populations. While population expansions have previously been detected using SNPs in global and European *A. thaliana* (The1001GenomesConsortium 2016; Lee, et al. 2017) and samples combined from multiple *B. distachyon* clonal groups (Minadakis, et al. 2023), our approach offers the advantage of identifying such signatures within a single population without requiring excessively deep sequencing or a large sample size. Additionally, it enables the inference of demographic history in studies that only generate methylation data and details the limits of inference when using methylation markers. Overall, our study highlights the value of methylation markers for understanding recent shifts in population size.

## Conclusions

Our findings demonstrate that CG methylation can provide new insights into population structure in clonal and selfing species, similar to its previously demonstrated utility in *A. thaliana* from America and the clonal seagrass *Zostera marina* (Yao, et al. 2023). We demonstrate the utility of methylation variants for uncovering very recent demographic events, where SNP markers are limited due to their slower rate of change. We also illustrate its usefulness in clonal groups, where the paucity of SNPs would render such inferences either impossible or necessitate very deep and costly levels of DNA sequencing. Conversely, at older timescales, methylation markers are unlikely to be informative because the high rates of change would have resulted in the violation of the infinite sites model. We show that SNPs and SMPs are complementary for generating demographic inferences over short and long timescales, highlighting the value of a recently developed method that leverages both sources of variation for inferring demography (Sellinger, et al. 2023). Our work demonstrates that methylomes contain unique evolutionary insights unattainable through studying genomic variation in isolation.

## Methods

### Genetic and methylation variant calling in Arabidopsis thaliana

We obtained the *A. thaliana* 1001 genomes (The1001GenomesConsortium 2016) SNP calls from https://1001genomes.org/data/GMI-MPI/releases/v3.1/. We only retained biallelic SNPs for our analyses. As C/T variant calls can become confounded with methylation calls for data generated using bisulfite sequencing, we generated sample-specific genome references for mapping bisulfite sequencing reads. We substituted the SNP calls for each sample into the TAIR10 genome reference (Lamesch, et al. 2012) using SAMtools (Danecek, et al. 2021), BCFtools consensus (Danecek, et al. 2021), and fastq split (Nassar, et al. 2023). We obtained bisulfite sequencing data for the 1001 *A. thaliana* from (Kawakatsu, et al. 2016). We first removed poor-quality reads using Trimmomatic (Bolger, et al. 2014) with the following options: ILLUMINACLIP:TruSeq3-SE.fa:2:30:10 LEADING:3 TRAILING:3 SLIDINGWINDOW:4:15 MINLEN:36 for single-end reads and ILLUMINACLIP:TruSeq3-PE.fa:2:30:10:2:True LEADING:3 TRAILING:3 SLIDINGWINDOW:4:20 MINLEN:36 for paired-end reads. We subsequently mapped reads to the sample-specific reference genomes using Bismark with the aid of the HISAT2 aligner (Krueger and Andrews 2011). We ran Bismark deduplicate_bismark to remove optical duplicates and ran bismark_methylation_extractor to extract methylation calls. Some samples were sequenced using both single-end and paired-end reads. For these samples, the cytosine counts were merged after methylation calling. We studied cytosine sites with a coverage of at least five reads similar to what was done in (Kawakatsu, et al. 2016). To identify the methylation state of a site, we performed a binomial test with the non-conversion rate being used as the hypothesized probability of success following the method in (Lister, et al. 2008). The non-conversion rate was calculated as the proportion of methylation in the chloroplast as done in (Lister, et al. 2008). Sites were assigned as methylated if the false discovery rate corrected *P*-values following the binomial test was less than 0.01. Otherwise, they were designated as unmethylated. We used jDMR (Hazarika, et al. 2022) to identify DMRs in the CG contexts as this method accounts for coverage when calculating likelihood scores for methylation states. For this, we called the methylation status per site using METHimpute (function: callMethylationSeparate) (Taudt, et al. 2018), extracted cytosine clusters in the CG context ensuring that there was a minimum of eight cytosines in each cluster (function: CfromFASTAv4 and makeReg with default options makeRegnull = c(FALSE), N.boot = 10>5, N.sim.C = “all”, fp.rate = 0.01, set.tol = 0.01), and identified the methylation states of the regions in each sample (function: runMethimputeRegions), ensuring that we used a minimum of eight cytosines and a coverage of three.

63 IBnr and 67 CEU samples overlapped between the SNP and methylation variant call files. We randomly subsampled 63 samples from the CEU population to ensure the numbers were comparable to those of the IBnr population. For our analyses, we only used sites where at least 80% of the samples within each population were genotyped. The proportions of variant SNP and SMP sites were calculated by dividing the number of sites with minor allele frequencies greater than 0.02 by the total number of variant and invariant sites.

### Population (epi)genetic metric estimation for Arabidopsis thaliana

To estimate π and Tajima’s *D*, we created fasta files for each genomic region containing the sequences for each sample. We substituted the variants from the SNP vcf into the TAIR10 reference genome using the approach described previously for each sample, extracted the region of interest, and converted it to a fasta format. The regions could be the clocklike regions described in (Yao, et al. 2023), genic regions comprising all exons, or non-coding regions comprising all regions other than exons. Genic regions were further split into gbM genes using the list for *A. thaliana* provided by (Williams, et al. 2023) and high SIFT-score (>0.75) and low SIFT-score (<0.3) genes using the list provided by (Hamala and Tiffin 2020).

We estimated π*_SNP_* and *D_SMP_* using pegas (Paradis 2010) and estimated π*_SMP_* and *D_SMP_* using the *D^m^* (Wang and Fan 2014). We only analyzed intervals that had at least three polymorphic sites and sequences with lengths greater than 10bp. We calculated the number of cytosines in each interval using METHimpute (Taudt, et al. 2018). To compare π and Tajima’s *D* among the different genomic contexts, we built a linear model where these metrics were the dependent variable, and the concatenated population and genomic context groupings were the predictor variables. We then applied a Tukey honest significant differences test using the agricolae R package (https://myaseen208.com/agricolae//index.html) to identify significant differences between individual predictor levels. We performed a correlation test to compare the π and Tajima’s *D* for SMPs and SNPs per exon.

We used VCFtools (Danecek, et al. 2011) to estimate the allele frequencies for SMPs and DMRs. We used PopLDdecay (Zhang, et al. 2019) to estimate pairwise LD using the r^2^ metric analyzing all sites with a minor allele frequency greater than 0.05. r^2^ values across every three base pair interval were averaged prior to plotting the decay curve.

We detected a total of 6,012 and 6,188 polymorphic noncoding STRs in the CEU and IBnr populations, respectively. To reduce computational time, we randomly sampled 100 STRs without replacement for each run and repeated this process 10 times for each population. We performed the sign test, the standardized difference test, and the Wilcoxon signed-rank test. The tests were conducted using the two-phase model (TPM) and the stepwise mutation model (SMM), as these models best describe mutational processes in STRs. The variance for TPM was set to 30, with 70% SMM in the TPM. We ran 1,000 iterations.

### Between population comparisons for Arabidopsis thaliana

We combined the CEU and IBnr vcfs and used VCF2Dis (https://github.com/BGI-shenzhen/VCF2Dis) to estimate the p-distance matrix. We used this matrix to generate neighbor-joining trees using FastME (Lefort, et al. 2015) and plotted it using ggtree (Yu, et al. 2017). We also performed DAPC to discriminate samples from the CEU and IBnr populations. We thinned the dataset so that only sites that were separated by 1Kbp were sampled to reduce the computational time. We converted vcf files into a genlight format using the *genomic_converter* function in the R package radiator (https://thierrygosselin.github.io/radiator/). We used the find.clusters function in adegenet (Jombart 2008) to identify the number of clusters in each dataset allowing for a maximum of 20 clusters for SMPs/DMRs and 30 clusters for SNPs. As DAPC regions require fully typed data, we imputed missing values using the mean value. Then, we performed cross-validation using adegenet (Jombart 2008) to identify the number of informative principal components. We retained a maximum of 15 principal components, used 90% of the data for training, and calculated group-wise assignment success. We centered but did not scale variables. We used 30 replicates for each principal component. Finally, we performed DAPC using adegenet (Jombart 2008).

### Methylation rate estimation *for Arabidopsis thaliana*

We generated the multihetsep files for SNPs for the same 5Mbp regions that were used for SMPs (described in the section immediately above). We combined the SNP and SMP multihetsep files. We randomly selected 20 samples from each population to ensure reasonable runtimes. We used the *Methylation_rate_estimation* function in SMCm to estimate forward and backward rates (Sellinger, et al. 2023). We set the mutation rate as 7×10⁻^9^ per site per generation and the recombination rate as 3.5×10⁻^8^ per site per generation. We ignored regional influences on methylation rates and estimated rates under a site-only model. We used the known estimates of forward (2.56×10⁻⁴ per generation per haploid methylome) and backward methylation rates (6.3×10⁻ ⁴ per generation per haploid methylome) for *A. thaliana* (van der Graaf, et al. 2015) as priors.

### Genetic and methylation variant calling in Brachypodium distachyon

We obtained whole genome sequencing data (29 samples) for *B. distachyon* from (Wilson, et al. 2019) and whole genome bisulfite sequencing data from (Eichten, et al. 2020). We used Bd21-3 v1.2 genome reference and the GenBank annotation from the Joint Genome Institute (http://phytozome.jgi.doe.gov/) for variant calling. We used Trimmomatic (Bolger, et al. 2014) to trim reads using the paired-end options as stated above for the whole genome sequencing data and the single-end option for the whole genome bisulfite sequencing data. We mapped whole genome sequencing data reads to the reference using bwa-mem (Li 2013). We used Picard Toolkit (https://broadinstitute.github.io/picard/) to add read groups and mark duplicate reads. We used SAMtools (Danecek, et al. 2021) to remove unpaired reads, remove reads with a mapping quality of less than 20, and sort reads. We called SNPs using GATK HaplotypeCaller (McKenna, et al. 2010). We only retained SNPs with depths greater than five and quality scores greater than 30 using BCFtools (Danecek, et al. 2021). Using this SNP variant call file, we perform methylation variant calling in the same way as was done for *A. thaliana*. We obtained the sample localities from (Filiz, et al. 2009) and created a map showing their occurrence with Map Maker (https://maps.co/).

We used a minor allele frequency threshold of 10% to identity polymorphic sites in *B. distachyon* due to the smaller number of samples being analyzed when compared to *A. thaliana*. For the analysis of SMPs and DMRs, we only used variants that occurred within CG-gbM regions identified in (Niederhuth, et al. 2016). We estimated mSFS, LD, p-distance matrices, π, and Tajima’s *D* in a similar way as we did for *A. thaliana*. We used the first principal component for the DAPC analyses.

### Generating the approximated transition matrix for demographic inference

We created a training and testing dataset consisting of a combined 60×10^3^ simulations (80%/20%, respectively). Of these 10×10^3^ were constant demographies (*i.e*., no demographic change) with a random population size drawn between 10×10^3^ and 200×10^3^, and the other 50×10^3^ simulations were random walk demographies, with time steps being defined for i ∈ {0,1,…,K − 1} using:

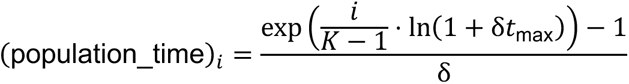

where *K*=42 is the number of time points, *δ* = 0.001 is the growth rate, and *t*_max_ = 2.1×10^6^ is the maximum time.

The population size was drawn for each of the *K*=42 time points in the following way: an initial population size was randomly selected between a minimum (*N_min* = 10×10^3^) and maximum (*N_max* = 1.2×10^6^) value. For each subsequent time point, a new population size was proposed by multiplying the previous size by a random factor between 0.1 and 10. If this proposed size fell outside the allowed range (N*_min* to *N_max*), new proposals were generated until an acceptable value was found. This process was repeated for all 42 time points, resulting in a sequence of population sizes that exhibit both gradual and abrupt changes while remaining within the specified bounds. Nevertheless, a sliding window average with a size of 6 and a step size of 1 was applied to smooth extreme changes. This approach for sampling demographies was adapted from (Sanchez, et al. 2021).

The obtained population sizes and times were passed onto *msprime* (Baumdicker, et al. 2022) to simulate the corresponding ancestral recombination graphs using a recombination rate of 3.4×10^−8^ per base pair and generation. For SNPs, branches were mutated at a rate of 7×10^−9^ per base pair and generation, leading to a *ρ* over *θ* ratio of 4.86.

Simulated and real data are summarized as a genotype matrix (*GM*) with dimensions *n* × *m*, where *n,m* ∈ N represent the number of samples and mutations, respectively. Each element *g_ij_* of *GM*, where i ∈ {1,…,n} and j ∈ {1,…,m}, denotes the presence (*g_ij_* = 1) or absence (*g_ij_* = 0) of mutation *j* in sample *i*. We generated a corresponding vector of positions of size *m* for each segregating site containing floating point numbers of the location in the genome.

For each random population size parameter, we used a genome of length 4Mbp,10 samples (ploidy=1), and the *GM* and positions vector to construct a density estimate of mutations along the genome (in theory, corresponding to the respective TMRCA at that region). To ensure the presence of enough markers, we segmented the genome into 20Kbp pieces and counted the number of heterozygous markers between all pairwise 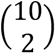 combinations. The 4Mbp sequences of all the comparisons were concatenated and log-transformed. Likewise, the population time steps were also log-scaled. To make sure that the number of markers per window fell approximately into the time windows, we scaled the markers by an arbitrarily chosen scaling factor (=*e*^8.2^) (Figure S12). The importance of the given scaling factor is negligible, due to all the transformation steps occurring in the pre-training, and therefore any noisy representation presented to the learning algorithm will be part of the training.

Each *msprime* simulation (under a random demographic history) yielded one *approximated* transition matrix, which was obtained by counting the transitions (scaled number of heterozygous sites in window t → window t+1) from one 20 Kbp window to the next using the following algorithm: a sliding window of size 2 and step size 1 was applied along the sequence and each pair of consecutive values in this window represents coordinates in the transition matrix, which was incremented at those positions leading to a filled matrix which remained unnormalized. Lastly, the entire transition matrix was incremented by one at each entry and scaled by the natural log to squeeze all values approximately into the same range to avoid training instability.

### Model description, training, and inference

The inference approach can be described as a supervised learning approach using the *approximated* transition matrices *x* with defined dimensions R*^L^*^=42×*K*=42^ as the input and the population size at each time step as the predicted variable. The input is a square matrix where both dimensions (K and L) are 42. The model architecture is an encoder-only transformer (Paszke, et al. 2019) (https://github.com/lucidrains/x-transformers). It consists of three main components: an input projection layer, a transformer encoder, and an output projection layer. It can be formalized as:

1. Input Projection: 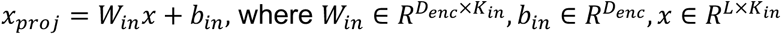
2. Transformer Encoder: 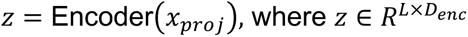 The Encoder employs a configuration with dimension *D_enc_* = 512, depth *N_enc_* = 12, number of attention heads *H_enc_* = 10 heads, Gated Linear Unit activation in feed-forward layers (Shazeer 2020), residual attention connections (He, et al. 2021), and rotary positional embeddings (Su, et al. 2024).
3. Dimension Reduction: 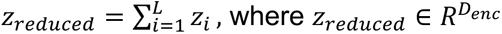
4. Output Projection: 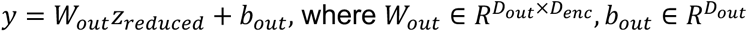

In this formalization, sequence length *L* = the input dimension of the transition matrix *K_in_* = 42, which represents the square input matrix of 42 x 42 dimensions, and the output dimension *D_out_* = 42.

The neural network model underwent training for 50 epochs, employing mean squared error as the loss function. The training process leverages the Accelerate framework from Hugging Face to enhance performance (Gugger, et al. 2022). An Accelerator object was configured with mixed precision using bfloat16 and gradient accumulation over 12 steps (with batch size 32). Throughout the training regiment, both the training and test losses were continuously monitored to evaluate model performance and assess convergence.

### Generating data for demographic inference

We first generated unique variant call files for each sample from the population variant call files using BCFtools (Danecek, et al. 2021). We then used the generate_multihetsep.py from MSMC-tools (https://github.com/stschiff/msmc-tools/tree/master) to combine all the samples per chromosome. For *A. thaliana* SNPs, we used the entire chromosome 1 as we needed a long contiguous fragment to generate the approximated transition matrix. For SMPs, we used 5Mbp segments in each of the five chromosomes (Table S3) as the SMP density was far higher. We used the list of genotyped cytosine positions as a genomic location mask for the SMPs. For *B. distachyon*, we included singletons to support the demographic inference given the small sample sizes in this species. The data were polarized by identifying the minor (less frequent) alleles and encoding them as derived, whereas the major alleles were encoded as ancestral. The resulting genotype matrix represents this polarized, unphased data, where each row corresponds to an individual and each column corresponds to a genomic position.

### Methylation simulation model

We explored the boundaries of our course-grained demographic inference by adding a new mutation model to msprime (Baumdicker, et al. 2022). We simulated a sawtooth demographic model with cyclical population size changes over time and a baseline effective population size of 2×10^5^ individuals. Simulations were conducted on a 5Mbp sequence with a recombination rate of 3.4×10^−8^ per base pair per generation (ploidy=1), to isolate the influence of methylation dynamics.

A custom matrix mutation simulation was implemented to model methylation and demethylation processes, employing 2 allelic states: 0 (unmethylated) and 1 (methylated). The model incorporates 3 key rates: the site introduction rate (*µ*), methylation rate (*β*), and demethylation rate (*δ*). The methylation and demethylation rates are measured as factors of the site entry rate, where *β* = the methylation factor·*µ* and *δ* = the demethylation factor·*µ*. The transition probabilities between the unmethylated and methylated states are determined by the ratio *β/*(*β*+*δ*) for methylation and *δ/*(*β*+*δ*) for demethylation. The resulting 2×2 transition matrix encapsulates these probabilities:

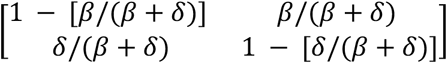

This framework allowed us to explore various methylation scenarios by adjusting *β* and *δ* relative to *µ*. We investigated cases where *β* = *µ* and *δ* = 2.5*µ*, *β* = 10*µ* and *δ* = 25*µ*, and *β* = 100*µ* and *δ* = 250*µ*, representing low, medium, and high methylation dynamics respectively, with the demethylation rate 2.5 times higher than the methylation rate. By varying the epimutation rate compared to the mutation rate, we test the robustness of our approach in correlating the number of markers in one Kbp windows with the average coalescent time. This proportionality, and the robustness of our method, is expected to decrease for higher epimutation rates, as these would violate the infinite site model for older coalescent times more strongly (due to homoplasy, (Vidalis, et al. 2016)).

## Declarations

### Availability of data and materials

The scripts for analyzing and plotting all the results, except for the demographic inference, have been published on GitHub (arunkumarramesh/methevolve) and archived using Zenodo (https://doi.org/10.5281/zenodo.14851293). The demographic inference method is available on GitHub (kevinkorfmann/aTMi) and has been archived using Zenodo (https://doi.org/10.5281/zenodo.13332833).

### Authors’ contributions

RA and AT designed the research. RA, KK, AW, BH, CG, and YW performed the research. RA, AT, KK, and AW wrote the paper. All authors read and approved the final manuscript.

## Acknowledgements

We would like to thank Thibaut Paul Patrick Sellinger for helping with the estimation of methylation rates. This work was supported by the de.NBI Cloud within the German Network for Bioinformatics Infrastructure (de.NBI) and ELIXIR-DE (Forschungszentrum Jülich and W-de.NBI-001, W-de.NBI-004, W-de.NBI-008, W-de.NBI-010, W-de.NBI-013, W-de.NBI-014, W-de.NBI-016, W-de.NBI-022).

