## Supplementary file for "Methylomes reveal recent evolutionary changes in populations of two plant species"

### Table of contents

Supplementary figure 1-12
Supplementary tables 1-3

### Supplementary figures

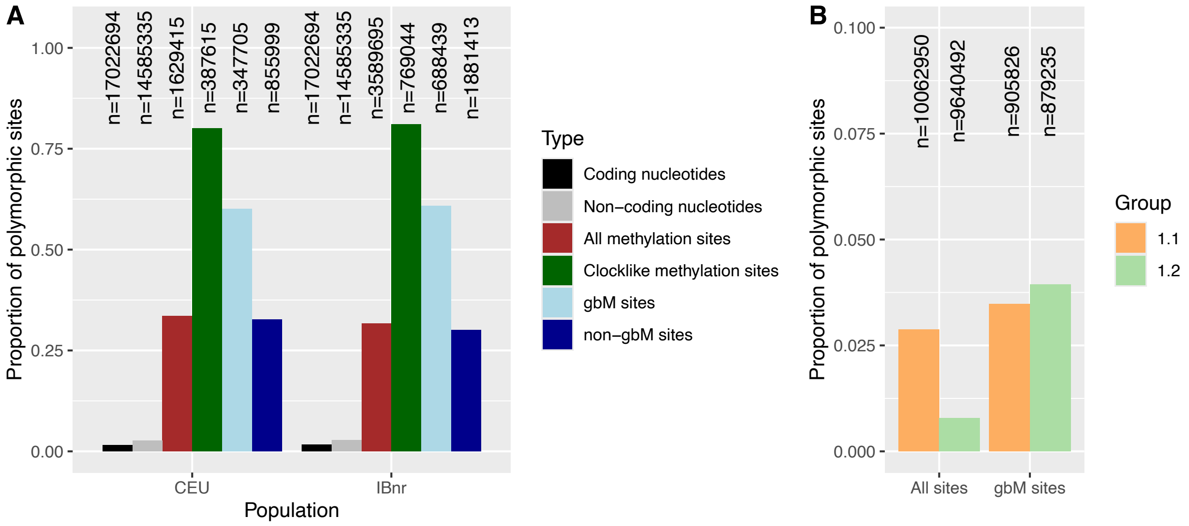


**Figure S1**. Proportion of polymorphic sites (**A**) Proportion of polymorphic nucleotide and CG methylation sites in the Central European (CEU) and the Iberian non-relicts (IBnr) *Arabidopsis thaliana* populations. (**B**) Proportion of polymorphic CG methylation sites for *Brachypodium distachyon* clonal groups 1.1 and 1.2. For (A) and (B), the numbers above the bars indicate the total number of variant and invariant sites per class.

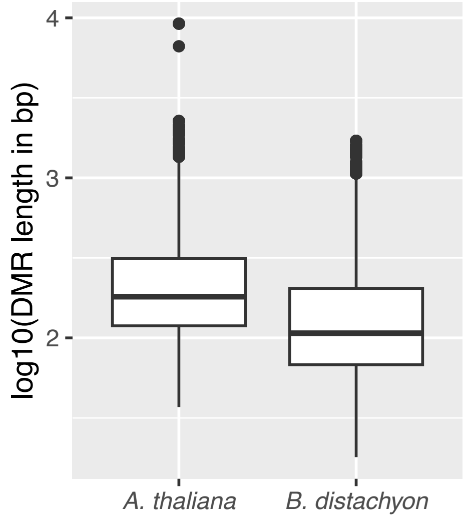


**Figure S2**. Lengths of differentially methylated regions (DMR) in *Arabidopsis thaliana* and *Brachypodium distachyon.*

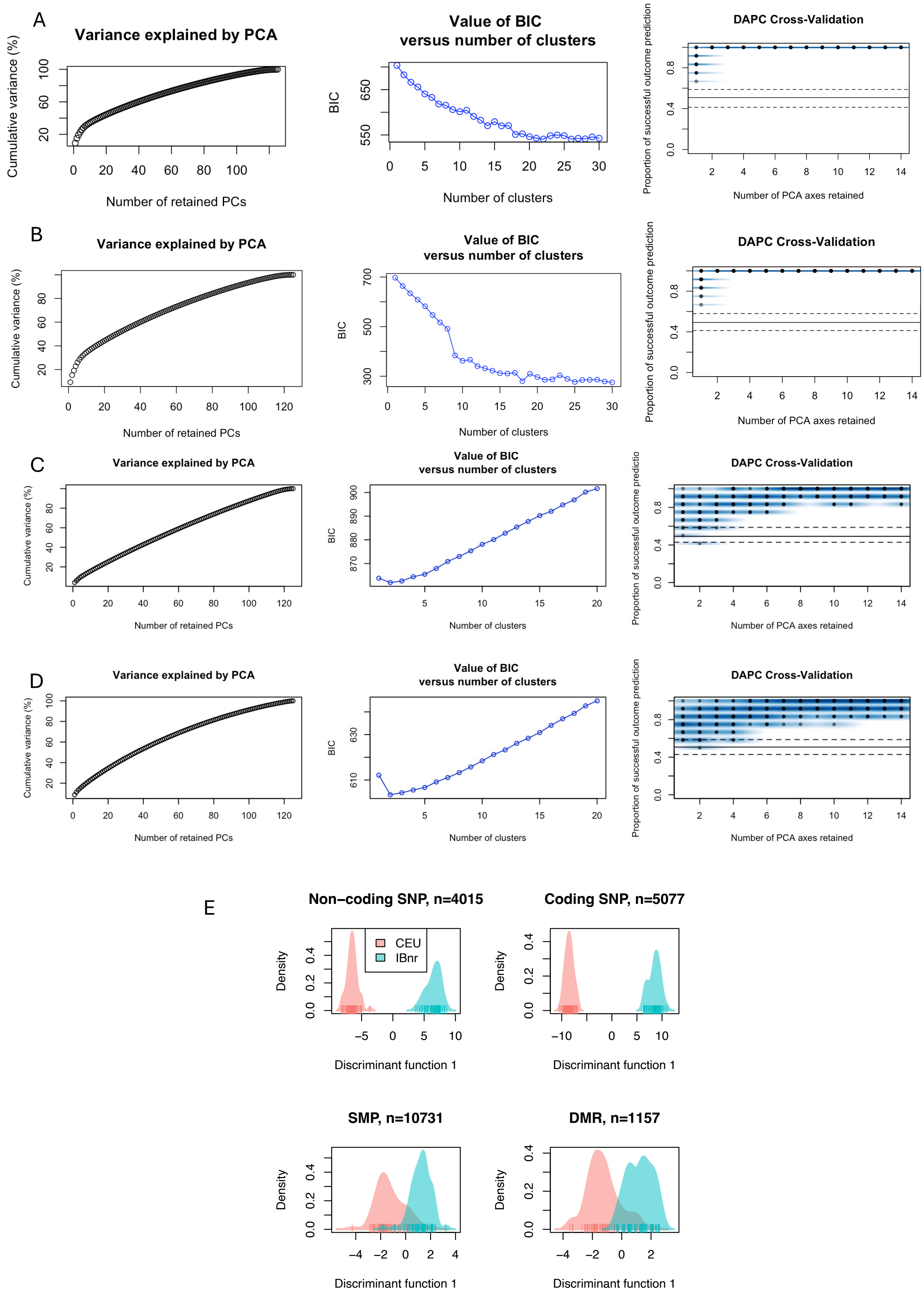


**Figure S3**. Cluster discrimination for single nucleotide polymorphisms (SNP), single methylation polymorphisms (SMP) and differentially methylated regions (DMR) datasets from the Central European (CEU) and the Iberian non-relicts (IBnr) *Arabidopsis thaliana*

populations. Clustering metrics for (**A**) Non-coding SNPs, (**B**) Coding SNPs, (**C**) Clocklike SMPs, and (**D**) Clocklike DMRs. For (A) – (D), each panel shows the cumulative variance explained by principal components (PC), Bayesian information criterion (BIC) versus the number of clusters, and the proportion of successful outcome predictions versus number of PCs used for the discriminant analysis of principal components analysis (DAPC). (**E**) DAPC for SNPs, SMPs, and DMRs in clocklike regions. Five principal components were retained for the DAPC. The numbers in the titles of each panel indicate the number of sites used for the analyses.

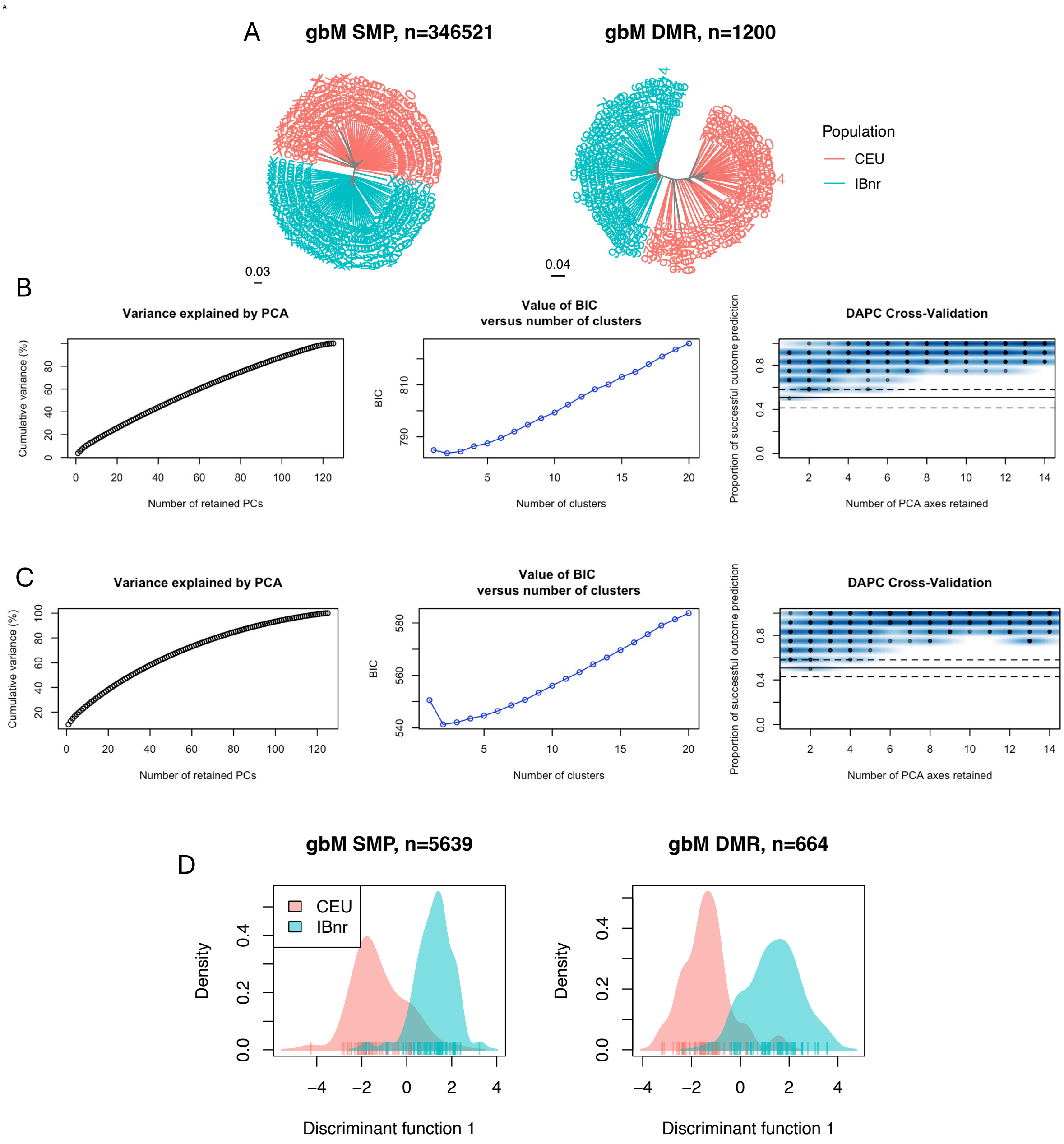


**Figure S4**. Using single methylation polymorphisms (SMP) and differentially methylated regions (DMR) that occur within gene body methylated genes (gbM) to differentiate the Central European (CEU) and the Iberian non-relicts (IBnr) *Arabidopsis thaliana* populations. (**A**) Neighbor-joining trees generated using p-distance matrices generated for SMPs and DMRs. Clustering metrics for (**B**) SMPs in gene body methylated (gbM) genes, and (**C**)

DMRs in gbM genes. For (B) and (C), each panel shows the cumulative variance explained by principal components (PC), Bayesian information criterion (BIC) versus the number of clusters, and the proportion of successful outcome predictions versus number of PCs used for the discriminant analysis of principal components analysis (DAPC). (**D**) DAPC for SMPs and DMRs. Five principal components were retained for the DAPC. The numbers in the titles of each panel indicates the number of sites used for the analyses.

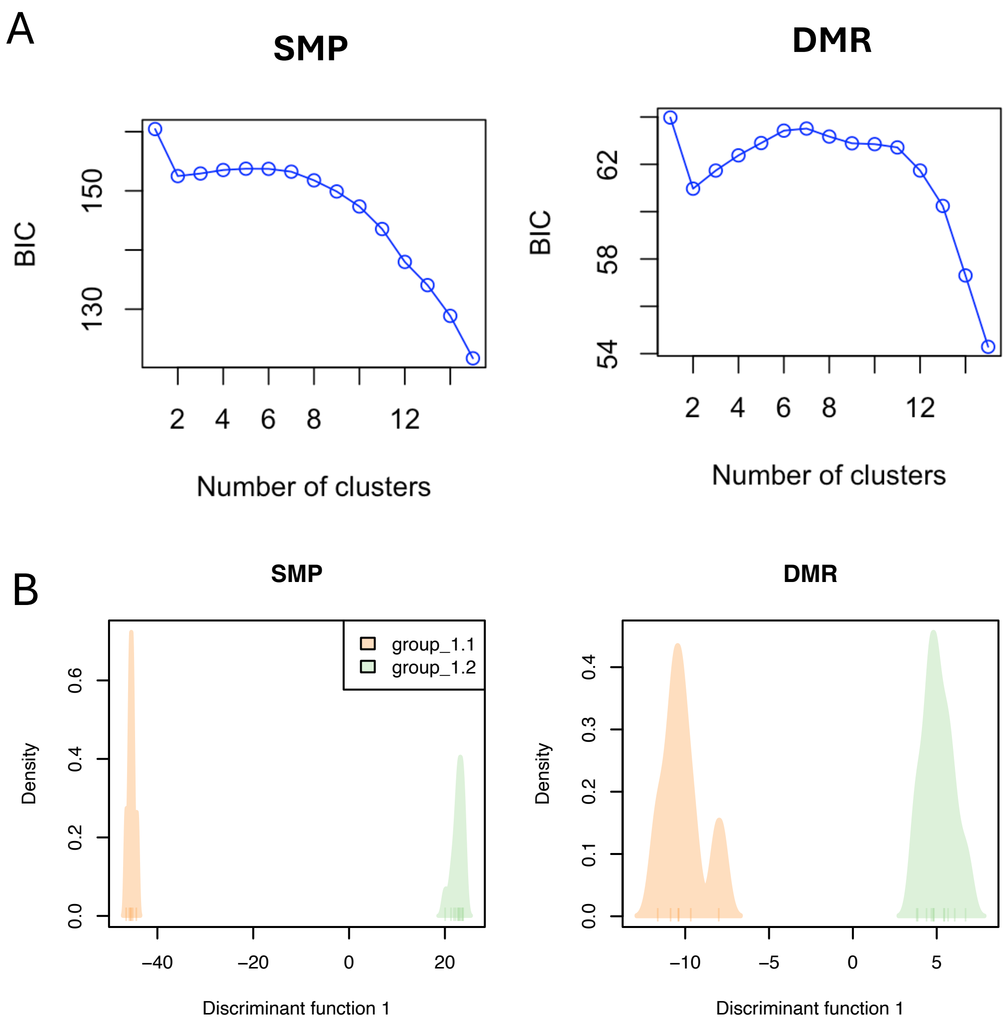


**Figure S5**. Subgroup discrimination for *Brachypodium distachyon*. (**A**) Bayesian information criterion (BIC) versus the number of clusters for single methylation polymorphisms (SMP) and differentially methylated regions (DMR). (**B**) Discriminant analysis of principal components analysis (DAPC) for SMPs and DMRs. 50422 SMP sites and 225 DMRs that occur within gene body methylated regions were used for both (A) and (B).

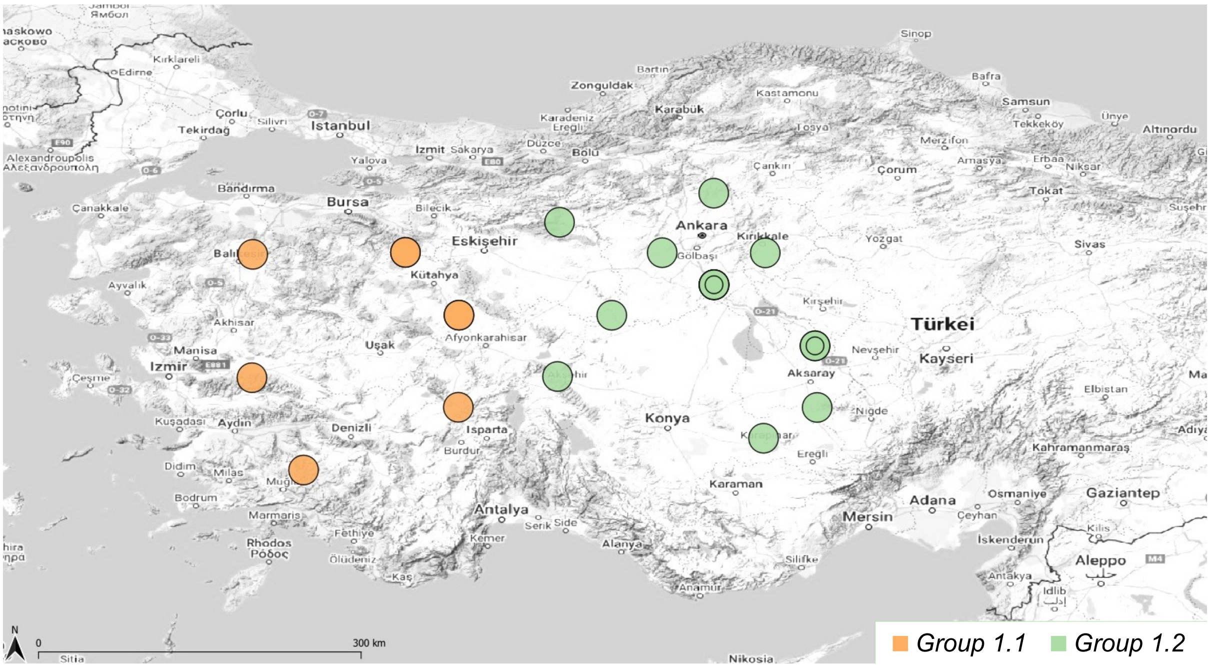


**Figure S6**. Longitudinal variation in the localities of *Brachypodium distachyon* samples from clonal groups 1.1 and 1.2. Circles within circles indicate multiple samples originating from the same site.

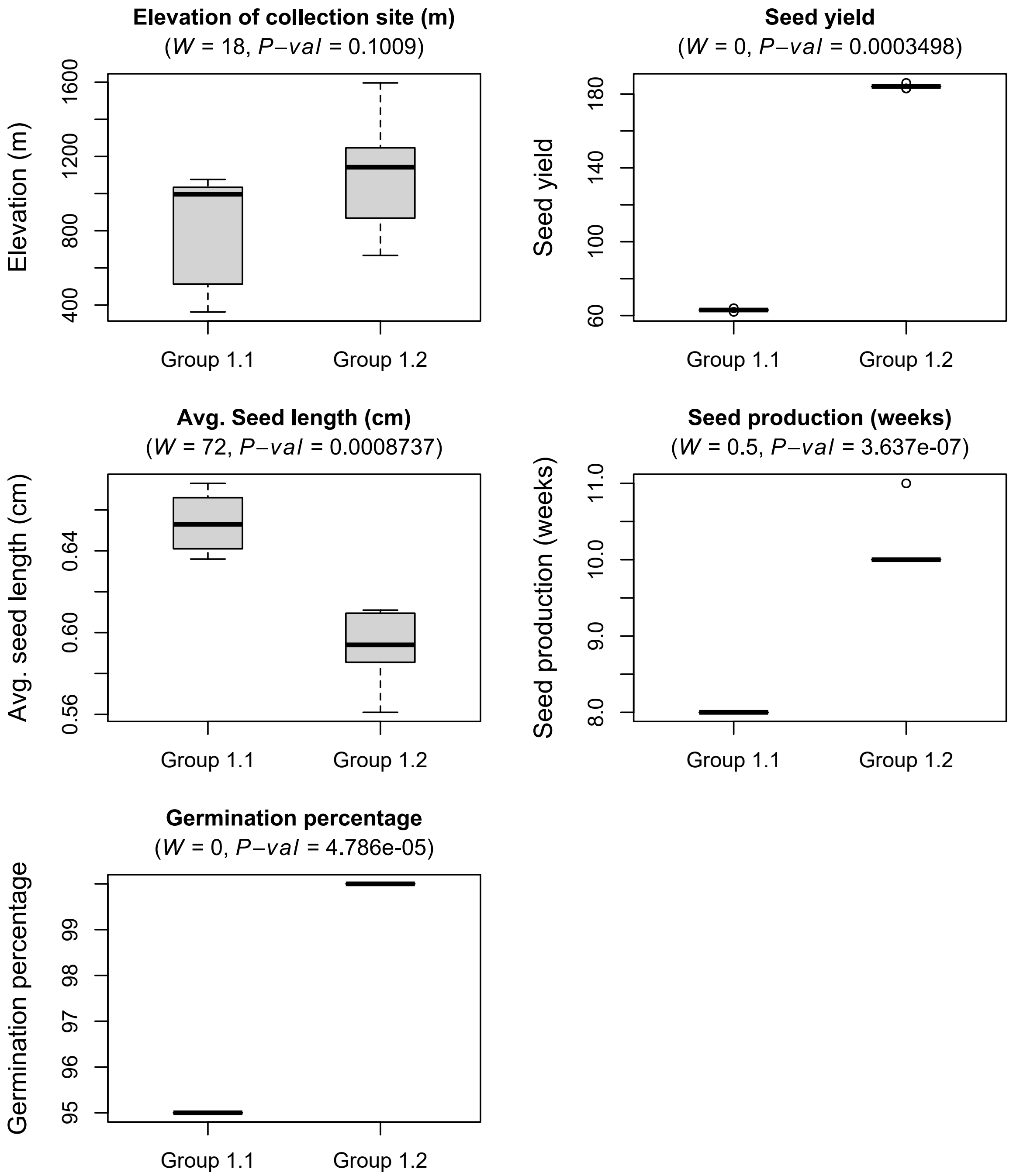


**Figure S7**. Phenotypic differences between samples from *Brachypodium distachyon* clonal groups 1.1 and 1.2. The *W* statistic and the *P*-values from Wilcoxon rank sum tests comparing the phenotypes from the two subgroups are indicated above the panels.

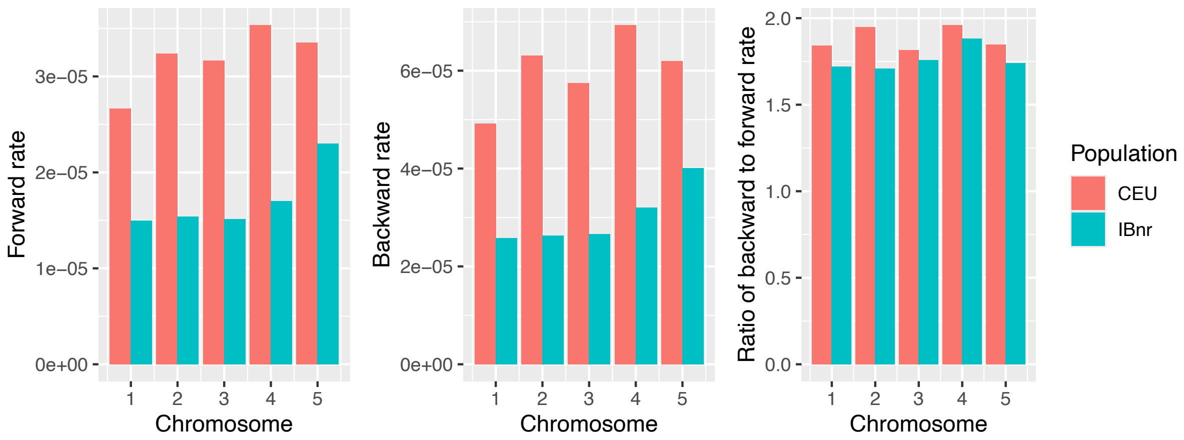


**Figure S8**. Forward and backward methylation rates for the central European (CEU) and Iberian non-relicts (IBnr) *Arabidopsis thaliana* populations.

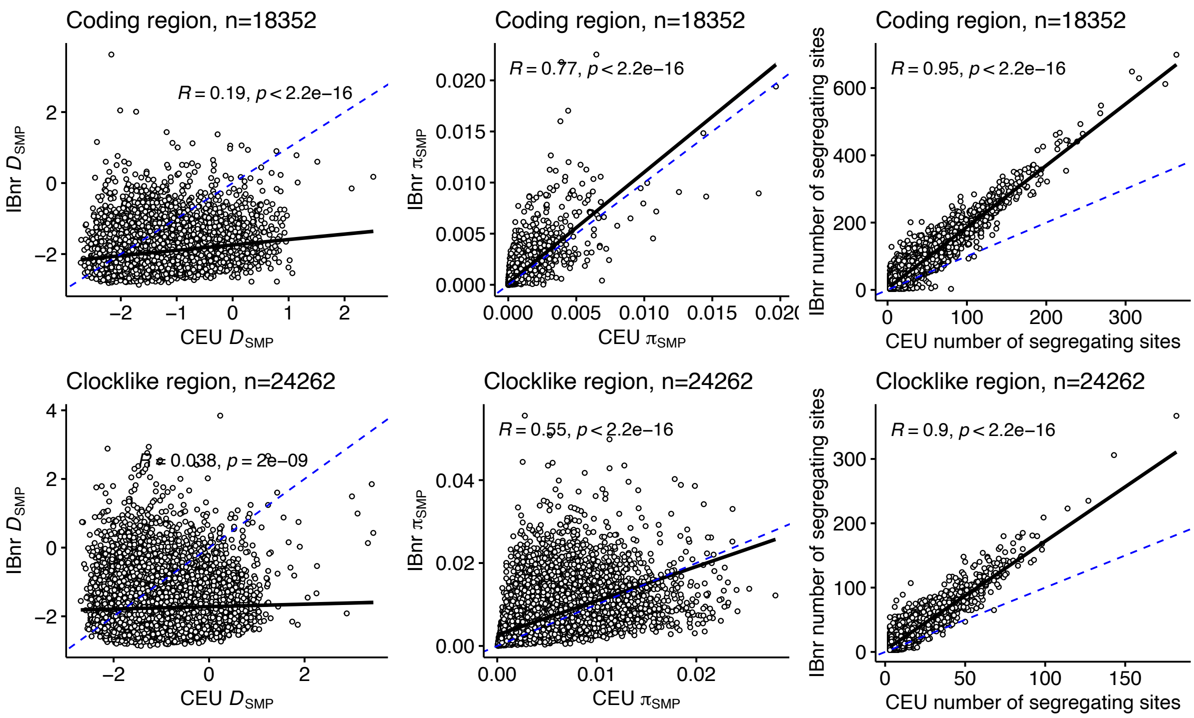


**Figure S9**. Comparison of $D_{SMP}$, π and the number of segregating sites for the *Arabidopsis thaliana* Central European (CEU) and Iberian non-relicts (IBnr) populations. The comparisons for exons within coding regions and for clocklike regions are shown. The black solid line is the linear regression fit. The blue dash line is the identity line. Correlation coefficients and *P*-values from the correlation test are shown within the panels. Number of regions are indicated in the panel titles. These values were used for generating Figure 3.

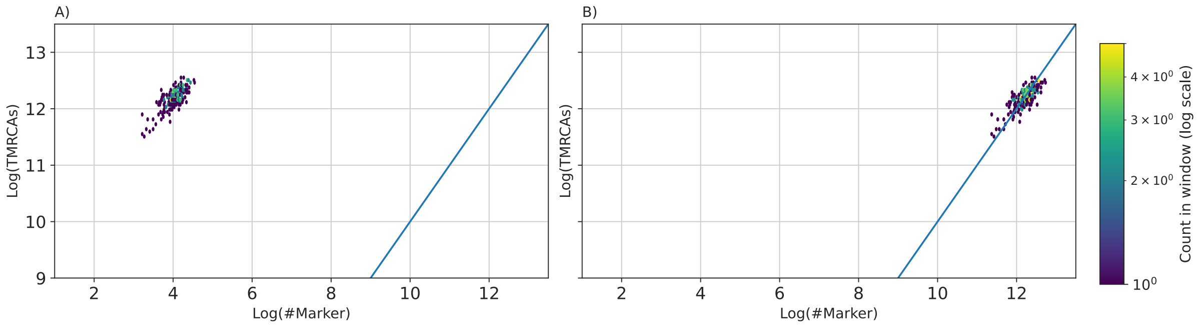


**Figure S10**. Empirical scaling factor justification. As mutations represent a sparse picture of underlying coalescing branches, they can also be regarded as noisy substitutes. Shown are correlations between the number of markers per 20Kbp window against the average time to the most recent common ancestor (TMRCA) in that window. A perfect correlation would be the straight line. However, shifting the line markers by scaling of $e^{8.2}$ corrects for the offset. The scaling factor has been visually chosen and does not influence the underlying result, as it was selected pre-training, and therefore only represents a data-transformation which is the same for all the training and testing samples.

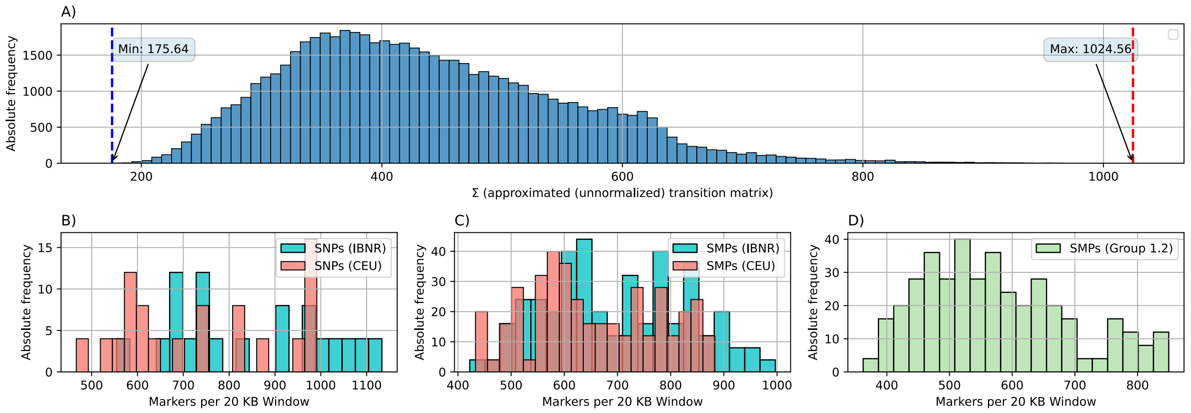


**Figure S11**. Matching distributions of single nucleotide polymorphisms (SNP) and single methylation polymorphisms (SMP) for in-distribution inference. (**A**) The distribution over the training and testing dataset presented by taking the sum of each entire (unnormalized) approximated transition matrix. The distribution of the number of markers for (**B**) *Arabidopsis thaliana* SNPs, (**C**) *A. thaliana* SMPs, and (**D**) *Brachypodium distachyon* SMPs. Distributions were generated for 20Kbp windows accumulated from 10 samples. Samples of sequence length between 1 and 3 Mbp have been drawn of the first chromosome for *A. thaliana* SNPs whereas samples have been drawn from chromosomes 1 to 5 for SMPs.

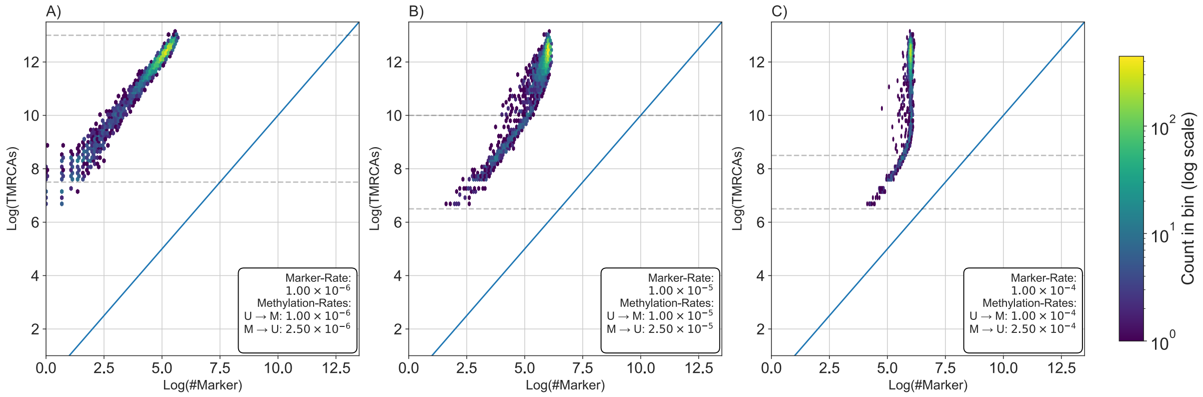


**Figure S12**. Simulating methylation markers with variable rates of change. We illustrate the effects of varying methylation and demethylation rates on the scaling property of the number of heterozygous markers to respective average time to the most recent common ancestor (TMRCA) per window. Methylation markers change at a rate of about (**A**) $10^{-6}$, (**B**) $10^{-5}$, or (**C**) $10^{-4}$ per site per generation. Backward methylation rate (methylated to unmethylated) is 2.5 times higher than the forward methylation rate (unmethylated to methylated). The dotted vertical lines indicate the timescales where the inference is likely to be informative.

### Supplementary tables

**Table S1**. List of samples used for this study.

| Accession | Group | Species |
|---|---|---|
| 6951 | CEU | *A. thaliana* |
| 8285 | CEU | *A. thaliana* |
| 6957 | CEU | *A. thaliana* |
| 8284 | CEU | *A. thaliana* |
| 9735 | CEU | *A. thaliana* |
| 428 | CEU | *A. thaliana* |
| 7207 | CEU | *A. thaliana* |
| 9973 | CEU | *A. thaliana* |
| 5921 | CEU | *A. thaliana* |
| 7520 | CEU | *A. thaliana* |
| 9696 | CEU | *A. thaliana* |
| 6445 | CEU | *A. thaliana* |
| 9731 | CEU | *A. thaliana* |
| 9694 | CEU | *A. thaliana* |
| 6956 | CEU | *A. thaliana* |
| 7120 | CEU | *A. thaliana* |
| 9728 | CEU | *A. thaliana* |
| 8365 | CEU | *A. thaliana* |

| | | |
|---|---|---|
| 9676 | CEU | *A. thaliana* |
| 6296 | CEU | *A. thaliana* |
| 424 | CEU | *A. thaliana* |
| 7350 | CEU | *A. thaliana* |
| 9727 | CEU | *A. thaliana* |
| 9679 | CEU | *A. thaliana* |
| 9668 | CEU | *A. thaliana* |
| 9729 | CEU | *A. thaliana* |
| 7424 | CEU | *A. thaliana* |
| 5950 | CEU | *A. thaliana* |
| 430 | CEU | *A. thaliana* |
| 9693 | CEU | *A. thaliana* |
| 7103 | CEU | *A. thaliana* |
| 6984 | CEU | *A. thaliana* |
| 8236 | CEU | *A. thaliana* |
| 6396 | CEU | *A. thaliana* |
| 7067 | CEU | *A. thaliana* |
| 9665 | CEU | *A. thaliana* |
| 5907 | CEU | *A. thaliana* |
| 7203 | CEU | *A. thaliana* |
| 10020 | CEU | *A. thaliana* |
| 9732 | CEU | *A. thaliana* |
| 5874 | CEU | *A. thaliana* |
| 6903 | CEU | *A. thaliana* |
| 7521 | CEU | *A. thaliana* |
| 6976 | CEU | *A. thaliana* |
| 9730 | CEU | *A. thaliana* |
| 9914 | CEU | *A. thaliana* |
| 9689 | CEU | *A. thaliana* |
| 6008 | CEU | *A. thaliana* |
| 8386 | CEU | *A. thaliana* |
| 5893 | CEU | *A. thaliana* |
| 403 | CEU | *A. thaliana* |
| 9684 | CEU | *A. thaliana* |
| 5993 | CEU | *A. thaliana* |
| 9915 | CEU | *A. thaliana* |
| 8290 | CEU | *A. thaliana* |
| 9671 | CEU | *A. thaliana* |
| 7296 | CEU | *A. thaliana* |
| 9733 | CEU | *A. thaliana* |
| 5890 | CEU | *A. thaliana* |
| 6390 | CEU | *A. thaliana* |

| | | |
|---|---|---|
| 410 | CEU | *A. thaliana* |
| 7177 | CEU | *A. thaliana* |
| 9690 | CEU | *A. thaliana* |
| 6933 | Ibnr | *A. thaliana* |
| 6970 | Ibnr | *A. thaliana* |
| 7328 | Ibnr | *A. thaliana* |
| 9507 | Ibnr | *A. thaliana* |
| 9510 | Ibnr | *A. thaliana* |
| 9511 | Ibnr | *A. thaliana* |
| 9514 | Ibnr | *A. thaliana* |
| 9515 | Ibnr | *A. thaliana* |
| 9521 | Ibnr | *A. thaliana* |
| 9522 | Ibnr | *A. thaliana* |
| 9524 | Ibnr | *A. thaliana* |
| 9525 | Ibnr | *A. thaliana* |
| 9534 | Ibnr | *A. thaliana* |
| 9535 | Ibnr | *A. thaliana* |
| 9537 | Ibnr | *A. thaliana* |
| 9540 | Ibnr | *A. thaliana* |
| 9541 | Ibnr | *A. thaliana* |
| 9544 | Ibnr | *A. thaliana* |
| 9547 | Ibnr | *A. thaliana* |
| 9556 | Ibnr | *A. thaliana* |
| 9557 | Ibnr | *A. thaliana* |
| 9560 | Ibnr | *A. thaliana* |
| 9562 | Ibnr | *A. thaliana* |
| 9564 | Ibnr | *A. thaliana* |
| 9567 | Ibnr | *A. thaliana* |
| 9568 | Ibnr | *A. thaliana* |
| 9577 | Ibnr | *A. thaliana* |
| 9582 | Ibnr | *A. thaliana* |
| 9588 | Ibnr | *A. thaliana* |
| 9594 | Ibnr | *A. thaliana* |
| 9821 | Ibnr | *A. thaliana* |
| 9822 | Ibnr | *A. thaliana* |
| 9825 | Ibnr | *A. thaliana* |
| 9833 | Ibnr | *A. thaliana* |
| 9834 | Ibnr | *A. thaliana* |
| 9836 | Ibnr | *A. thaliana* |
| 9840 | Ibnr | *A. thaliana* |
| 9841 | Ibnr | *A. thaliana* |
| 9843 | Ibnr | *A. thaliana* |

| | | |
|---|---|---|
| 9844 | Ibnr | *A. thaliana* |
| 9845 | Ibnr | *A. thaliana* |
| 9852 | Ibnr | *A. thaliana* |
| 9856 | Ibnr | *A. thaliana* |
| 9857 | Ibnr | *A. thaliana* |
| 9859 | Ibnr | *A. thaliana* |
| 9861 | Ibnr | *A. thaliana* |
| 9864 | Ibnr | *A. thaliana* |
| 9867 | Ibnr | *A. thaliana* |
| 9868 | Ibnr | *A. thaliana* |
| 9870 | Ibnr | *A. thaliana* |
| 9873 | Ibnr | *A. thaliana* |
| 9876 | Ibnr | *A. thaliana* |
| 9878 | Ibnr | *A. thaliana* |
| 9886 | Ibnr | *A. thaliana* |
| 9888 | Ibnr | *A. thaliana* |
| 9899 | Ibnr | *A. thaliana* |
| 9900 | Ibnr | *A. thaliana* |
| 9902 | Ibnr | *A. thaliana* |
| 9904 | Ibnr | *A. thaliana* |
| 9946 | Ibnr | *A. thaliana* |
| 9950 | Ibnr | *A. thaliana* |
| 6971 | Ibnr | *A. thaliana* |
| 7327 | Ibnr | *A. thaliana* |
| BdTR1e | 1.1 | *B. distachyon* |
| BdTR1h | 1.1 | *B. distachyon* |
| BdTR1j | 1.1 | *B. distachyon* |
| BdTR1k | 1.1 | *B. distachyon* |
| BdTR1m | 1.1 | *B. distachyon* |
| BdTR1n | 1.1 | *B. distachyon* |
| BdTR2b | 1.2 | *B. distachyon* |
| BdTR2c | 1.2 | *B. distachyon* |
| BdTR2d | 1.2 | *B. distachyon* |
| BdTR2g | 1.2 | *B. distachyon* |
| BdTR2h | 1.2 | *B. distachyon* |
| BdTR2j | 1.2 | *B. distachyon* |
| BdTR2k | 1.2 | *B. distachyon* |
| BdTR2m | 1.2 | *B. distachyon* |
| BdTR2n | 1.2 | *B. distachyon* |
| BdTR2p | 1.2 | *B. distachyon* |
| BdTR2r | 1.2 | *B. distachyon* |
| BdTR2s | 1.2 | *B. distachyon* |

**Table S2**. Tests for heterozygosity deficit under the two-phase model (T.P.M.) and stepwise mutation model (S.M.M.) for short tandem repeats (STR) from the Central European (CEU) and the Iberian non-relicts (IBnr) *Arabidopsis thaliana* populations. 100 STRs were randomly sampled from the entire STR dataset for each replicate.

| Population | Replicate | Mutation model | Sign test | | | | Standardised difference test | | Wilcoxon test | | |
|---|---|---|---|---|---|---|---|---|---|---|---|
| | | | Expected number of loci with heterozygosity excess | Number of loci with heterozygosity deficiency | Number of loci with heterozygosity excess | P-value | T2 | P-value | Heterozygosity deficit (one-tail P-value) | Heterozygosity excess (one-tail P-value) | Heterozygosity deficit/excess (two-tail P-value) |
| **CEU** | 1 | T.P.M. | 57.96 | 70 | 30 | <0.00001* | -7.67 | <0.00001* | <0.00001* | 1 | <0.00001* |
| **CEU** | 1 | S.M.M. | 58.56 | 86 | 14 | <0.00001* | -16.2 | <0.00001* | <0.00001* | 1 | <0.00001* |
| **CEU** | 2 | T.P.M. | 58.26 | 73 | 27 | <0.00001* | -9.32 | <0.00001* | <0.00001* | 1 | <0.00001* |
| **CEU** | 2 | S.M.M. | 58.65 | 87 | 13 | <0.00001* | -17.5 | <0.00001* | <0.00001* | 1 | <0.00001* |
| **CEU** | 3 | T.P.M. | 57.99 | 63 | 37 | 0.00002* | -6.72 | <0.00001* | 0.00001* | 0.99999 | <0.00001* |
| **CEU** | 3 | S.M.M. | 58.46 | 77 | 23 | <0.00001* | -14.2 | <0.00001* | <0.00001* | 1 | <0.00001* |
| **CEU** | 4 | T.P.M. | 57.68 | 75 | 25 | <0.00001* | -9.26 | <0.00001* | <0.00001* | 1 | <0.00001* |
| **CEU** | 4 | S.M.M. | 58.49 | 89 | 21 | <0.00001* | -17.6 | <0.00001* | <0.00001* | 1 | <0.00001* |
| **CEU** | 5 | T.P.M. | 58.13 | 66 | 34 | <0.00001* | -9.02 | <0.00001* | <0.00001* | 1 | <0.00001* |
| **CEU** | 5 | S.M.M. | 58.34 | 80 | 20 | <0.00001* | -17.6 | <0.00001* | <0.00001* | 1 | <0.00001* |
| **CEU** | 6 | T.P.M. | 57.22 | 65 | 35 | <0.00001* | -6.65 | <0.00001* | <0.00001* | 1 | <0.00001* |
| **CEU** | 6 | S.M.M. | 57.82 | 81 | 19 | <0.00001* | -14.1 | <0.00001* | <0.00001* | 1 | <0.00001* |
| **CEU** | 7 | T.P.M. | 57.57 | 64 | 36 | 0.00001* | -6.63 | <0.00001* | <0.00001* | 1 | <0.00001* |
| **CEU** | 7 | S.M.M. | 58.02 | 84 | 16 | <0.00001* | -14.2 | <0.00001* | <0.00001* | 1 | <0.00001* |
| **CEU** | 8 | T.P.M. | 57.93 | 71 | 29 | <0.00001* | -8.24 | <0.00001* | <0.00001* | 1 | <0.00001* |
| **CEU** | 8 | S.M.M. | 58.6 | 84 | 16 | <0.00001* | -16.6 | <0.00001* | <0.00001* | 1 | <0.00001* |
| **CEU** | 9 | T.P.M. | 57.91 | 73 | 27 | <0.00001* | -6.71 | <0.00001* | <0.00001* | 1 | <0.00001* |
| **CEU** | 9 | S.M.M. | 58.46 | 86 | 14 | <0.00001* | -14.3 | <0.00001* | <0.00001* | 1 | <0.00001* |
| **CEU** | 10 | T.P.M. | 58.33 | 65 | 35 | <0.00001* | -8.09 | <0.00001* | <0.00001* | 1 | <0.00001* |
| **CEU** | 10 | S.M.M. | 58.42 | 81 | 19 | <0.00001* | -16.8 | <0.00001* | <0.00001* | 1 | <0.00001* |
| **Ibnr** | 1 | T.P.M. | 57.46 | 70 | 30 | <0.00001* | -7.54 | <0.00001* | <0.00001* | 1 | <0.00001* |
| **Ibnr** | 1 | S.M.M. | 57.88 | 86 | 14 | <0.00001* | -17.2 | <0.00001* | <0.00001* | 1 | <0.00001* |
| **Ibnr** | 2 | T.P.M. | 58.29 | 69 | 31 | <0.00001* | -8.61 | <0.00001* | <0.00001* | 1 | <0.00001* |
| **Ibnr** | 2 | S.M.M. | 58.49 | 84 | 16 | <0.00001* | -20.1 | <0.00001* | <0.00001* | 1 | <0.00001* |
| **Ibnr** | 3 | T.P.M. | 57.78 | 60 | 40 | 0.00024* | -3.33 | 0.00044* | 0.00107* | 0.99895 | 0.00213* |
| **Ibnr** | 3 | S.M.M. | 57.89 | 76 | 24 | <0.00001* | -11.3 | <0.00001* | <0.00001* | 1 | <0.00001* |
| **Ibnr** | 4 | T.P.M. | 57.51 | 67 | 32 | <0.00001* | -8.02 | <0.00001* | <0.00001* | 1 | <0.00001* |
| **Ibnr** | 4 | S.M.M. | 57.88 | 89 | 10 | <0.00001* | -17.7 | <0.00001* | <0.00001* | 1 | <0.00001* |
| **Ibnr** | 5 | T.P.M. | 57.79 | 65 | 35 | <0.00001* | -8.03 | <0.00001* | <0.00001* | 1 | <0.00001* |
| **Ibnr** | 5 | S.M.M. | 58.31 | 81 | 19 | <0.00001* | -18.5 | <0.00001* | <0.00001* | 1 | <0.00001* |
| **Ibnr** | 6 | T.P.M. | 57.88 | 67 | 33 | <0.00001* | -7.84 | <0.00001* | <0.00001* | 1 | <0.00001* |
| **Ibnr** | 6 | S.M.M. | 58.05 | 86 | 14 | <0.00001* | -17.8 | <0.00001* | <0.00001* | 1 | <0.00001* |
| **Ibnr** | 7 | T.P.M. | 57.95 | 68 | 32 | <0.00001* | -5.48 | <0.00001* | <0.00001* | 1 | <0.00001* |
| **Ibnr** | 7 | S.M.M. | 58.15 | 91 | 9 | <0.00001* | -14 | <0.00001* | <0.00001* | 1 | <0.00001* |
| **Ibnr** | 8 | T.P.M. | 57.84 | 65 | 34 | <0.00001* | -7.78 | <0.00001* | 0.00001* | 1 | 0.00001* |
| **Ibnr** | 8 | S.M.M. | 58.2 | 83 | 16 | <0.00001* | -17.9 | <0.00001* | <0.00001* | 1 | <0.00001* |
| **Ibnr** | 9 | T.P.M. | 58.27 | 58 | 42 | 0.00073* | -6.95 | <0.00001* | 0.00004* | 0.99996 | 0.00008* |
| **Ibnr** | 9 | S.M.M. | 58.29 | 79 | 21 | <0.00001* | -17.1 | <0.00001* | <0.00001* | 1 | <0.00001* |
| **Ibnr** | 10 | T.P.M. | 57.37 | 64 | 36 | 0.00001* | -7.24 | <0.00001* | <0.00001* | 1 | <0.00001* |
| **Ibnr** | 10 | S.M.M. | 57.64 | 83 | 17 | <0.00001* | -17.3 | <0.00001* | <0.00001* | 1 | <0.00001* |

**Table S3**. Genomic intervals used for demographic inference using methylation variants.

| Species | Chromosome | Start | End |
|---|---|---|---|
| *A. thaliana* | 1 | 5500000 | 8500000 |
| *A. thaliana* | 2 | 15000000 | 18000000 |
| *A. thaliana* | 3 | 5000000 | 8000000 |
| *A. thaliana* | 4 | 12000000 | 15000000 |
| *A. thaliana* | 5 | 500000 | 3500000 |
| *B. distachyon* | 1 | 4000000 | 9000000 |
| *B. distachyon* | 2 | 10000000 | 15000000 |
| *B. distachyon* | 3 | 4000000 | 9000000 |
| *B. distachyon* | 4 | 10000000 | 15000000 |
| *B. distachyon* | 5 | 10000000 | 15000000 |
